# scMultiNODE: Recovering Developmental Dynamics in Weakly Resolved Modalities through Multi-Modal Integration

**DOI:** 10.1101/2024.10.27.620531

**Authors:** Jiaqi Zhang, Manav Chakravarthy, Ritambhara Singh

## Abstract

Measuring multiple genomic profiles (i.e., modalities) over time enhances our understanding of cell development. However, these modalities are not equally analyzable. Sparse and poorly separated measurements, such as chromatin accessibility, are difficult to interpret on their own, and the limited availability of temporal single-cell co-assay data leaves such modalities without the cross-modal cell correspondence needed to borrow structure from richer ones. As a result, critical developmental analyses and cross-modal comparisons remain underperformed for these weak modalities. Additionally, existing methods cannot integrate separately sequenced measurements while preserving cellular dynamics. We present scMultiNODE, an integration model that combines gene expression and chromatin accessibility measurements from multiple discrete timepoints for single cells, without relying on any prior cell-to-cell correspondence across modalities or time. scMultiNODE employs a scalable Quantized Gromov-Wasserstein optimal transport method to align cells across measurements, and neural ordinary differential equations with a dynamic regularization term to model cell development in a shared latent space, capturing cellular dynamics while preserving cell-type variations. Experiments across four developmental single-cell datasets demonstrate that scMultiNODE integrates separately sequenced measurements over time more effectively than existing methods that overlook cellular dynamics. Crucially, by anchoring chromatin accessibility to the shared dynamical latent space, scMultiNODE recovers developmental trajectories and cell annotations from scATAC-seq that this modality cannot support in isolation. The resulting joint latent space further supports important multi-modal downstream analyses, including complex trajectory investigation and cross-modal label transfer for cell annotation. The data, code, and supplementary notes are publicly available at https://github.com/rsinghlab/scMultiNODE.

## 1. Introduction

Biological systems are inherently dynamic, constantly changing and adapting over time, and function at different scales, like organisms, cells, and molecules. Analyzing dynamic processes at the cell level reveals how cells grow, divide, and differentiate into various cell types.^1,2^ Understanding these cellular dynamics enhances our understanding of cell development and disease progression. Advances in single-cell measurements taken over time support this study by capturing high-resolution genomic snapshots of different cell states. Recent research increasingly involves temporally profiling multiple developmental single-cell measurements (or modalities), uncovering heterogeneity within the same tissue type and cellular dynamics across developmental stages.^3,4^ For instance, Fleck et al.^5^ collected gene expression and chromatin accessibility profiles across a developmental time course of brain organoids, capturing key processes such as neurogenesis. These modalities, however, are not equally informative. They differ not only in the biological signal they capture but in how readily that signal can be analyzed, and this asymmetry shapes what can be learned from each alone. In this paper, we present a deep learning model that integrates gene expression and chromatin accessibility modalities across discrete single-cell timepoints, enabling the characterization of cellular dynamics.

While existing studies^6–9^ have provided valuable insights on modeling cellular development using either gene expression or chromatin accessibility, each modality offers only a partial and unevenly reliable view. Gene expression provides detailed transcriptional readouts but is affected by dropouts and mRNA degradation,^10^ whereas chromatin accessibility reports regulatory potential yet is sparse, high-dimensional, and of low specificity, leaving it difficult to interpret in isolation and typically lacking cell-type annotations.^11^ This reflects not just a difference in content, but in analyzability. Core developmental analyses, such as reconstructing trajectories, ordering cells in pseudotime, and assigning cell identities, are unreliable or altogether infeasible when performed on a sparse, weakly separated modality such as chromatin accessibility on its own, because its cells do not cleanly resolve into the groups and paths these analyses depend on. Integration is usually motivated by completeness, such that combining complementary modalities yields a fuller description of a biological system.^12,13^ We emphasize a second, less appreciated consequence. Because the modalities share underlying biology, integration offers the opportunity to transfer the analytical strength of a richer modality to a weaker one, anchoring the poorly resolved modality within a shared representation so that analyses it cannot support alone become possible. Realizing this opportunity, rather than merely fusing modalities, is a central aim of our work.

In practice, however, the structure of the data works against this. Single-cell sequencing is inherently destructive, so each modality is typically measured in different cell populations, yielding datasets that are “unaligned”, where no cell-to-cell correspondence exists between modalities. Co-assay technologies that profile multiple modalities within the same cells (e.g., gene expression and chromatin accessibility)^14,15^ do provide such correspondence, but they remain resource-intensive and are often limited by low sequencing depth,^16,17^ which restricts their scalability across developmental timepoints and complex tissues. The unaligned setting sharpens the asymmetry described above. Without measured correspondence, there is no direct bridge along which a weak modality can borrow structure from a strong one, and the weak modality frequently lacks annotations altogether or carries only sparse, biased ones that are difficult to use. Consequently, most temporally resolved multi-modal datasets remain *unaligned across modalities and time*, so the analyses a weak modality would benefit from most, like trajectory inference and cross-modal comparison, are the hardest to perform on it.

Currently, there are no computational methods that can jointly integrate unaligned multimodal temporal datasets and explicitly model cellular dynamics over time. Existing approaches, primarily applied to static snapshot datasets, either align different modalities by identifying correlated patterns or common biological signals across modalities^19,20^ or adopt alignment techniques that better capture intricate cross-modality structures.^18,21–23^ They achieve flexible integration, particularly by aligning cell types across modalities, but treat integration as a symmetric fusion aimed at completeness and do not model how cells evolve over time. Although the recent scTIE model^24^ combines multiple timepoints from temporal data into a shared space for gene regulation analysis, it is not applicable to unaligned datasets and does not explicitly model cellular development. As a result, the integrated latent spaces of existing methods do not naturally encode cellular dynamics,^25^ limiting their ability to resolve temporally dependent transitions and branching developmental trajectories. Fig. 1B and Sec. 4.2 show state-of-the-art integration methods (e.g., SCOTv2^18^) learning an alignment of developmental multi-modal single-cell data but yielding trajectories that break continuity, contradict reference lineages, or connect unrelated cell groups, which has a direct and overlooked cost for weak modalities. A method that cannot represent developmental structure at all cannot transfer that structure to a poorly resolved modality, so even where static labels can be shared, dynamic analyses on the weak modality remain out of reach. Closing this gap requires a method that integrates unaligned temporal multi-modal data, explicitly models cellular dynamics, and thereby carries developmental structure to the modality that cannot recover it alone.

**Fig. 1.**
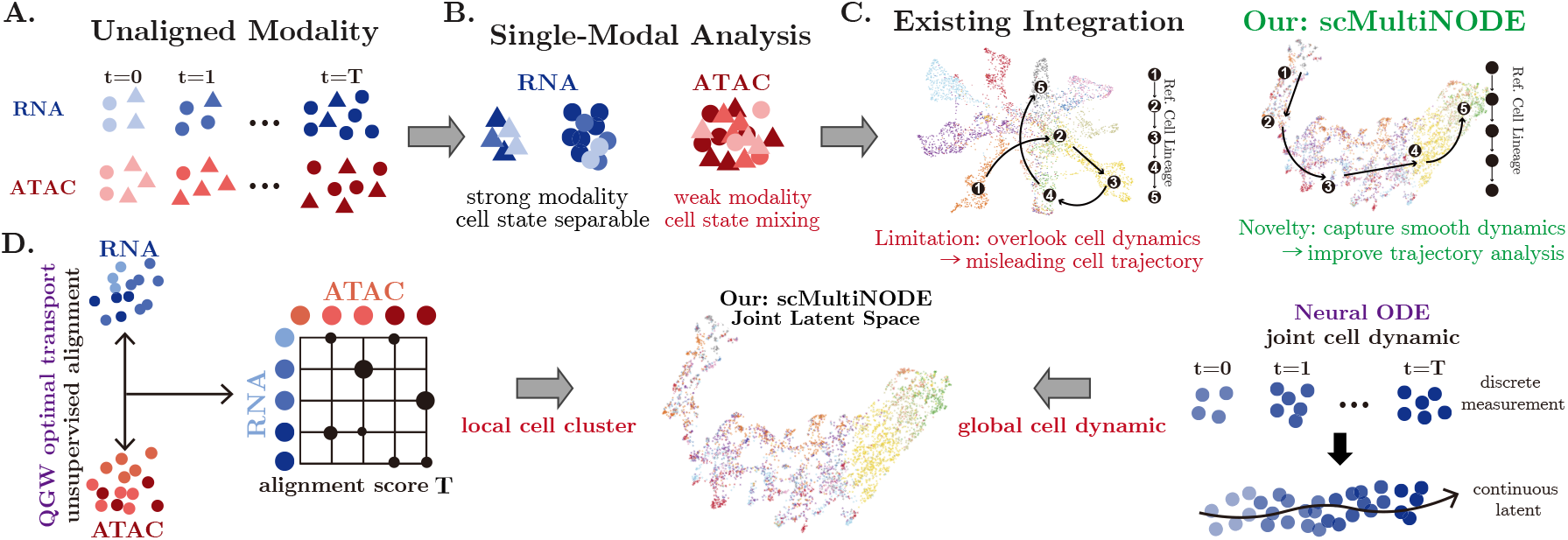
Model overview. (**A**) We integrate multi-timepoint scRNA-seq (gene expression) and scATAC-seq (chromatin accessibility) data without prior cell-to-cell correspondences across modalities or timepoints. (**B**) Single-modal analysis on the weaker modality limits accurate cell-state identification. (**C**) Existing integration methods (e.g., SCOTv2^18^) improve the weaker modality by leveraging the stronger one, but often fail to capture continuous cellular dynamics, yielding trajectories (solid arrows) that break continuity, contradict reference lineages, or connect unrelated cell groups. In contrast, our `scMultiNODE` explicitly models smooth and continuous dynamics, improving temporal analysis. (**D**) `scMultiNODE` aligns multi-modal datasets using QGW optimal transport to infer cross-modal cell correspondences while preserving cell-type structure, and models continuous cell dynamics with neural ODEs.

We propose single-cell Multi-modal Neural Ordinary Differential Equation (`scMultiNODE`), which integrates gene expression (scRNA-seq) and chromatin accessibility (scATAC-seq) profiles while explicitly modeling cellular dynamics over multiple timepoints (Fig. 1). `scMultiNODE` is a novel model that does not require prior cell-to-cell correspondence information across modalities or timepoints, as opposed to other multi-modal dynamic modeling approaches. Specifically, `scMultiNODE` aligns cells across modalities and timepoints using Quantized Gromov-Wasserstein (QGW) optimal transport,^26^ which identifies biologically similar cells and scales to large-scale datasets. Guided by this alignment, `scMultiNODE` constructs a joint latent space and explicitly learns smooth developmental trajectories using a single shared neural ordinary differential equation (ODE). Critically, because both modalities are governed by the same learned dynamics in this joint space, a sparse and weakly separated modality such as chromatin accessibility is placed onto the developmental flow inferred from the cleaner modality, rather than being modeled in isolation. As a result, `scMultiNODE` is the first method to preserve both local cell relationships (e.g., cell type distinctions) and global cellular dynamics (e.g., complex developmental trajectories) across multiple modalities during integration (Fig. 1D), a capability not achieved by previous multi-modal integration methods.

We evaluate `scMultiNODE` on four developmental single-cell datasets, comprising two coassay datasets and four unaligned datasets, using scRNA-seq and scATAC-seq assays measured at multiple timepoints across different species and tissues. Our analyses demonstrate that `scMultiNODE` integrates gene expression and chromatin accessibility measurements well in both unaligned and co-assay datasets. Moreover, `scMultiNODE` significantly outperforms baseline integrative models in capturing cellular dynamics while retaining cell-type variation. Beyond achieving high-quality integration, we demonstrate that `scMultiNODE` performs corrective integration. By leveraging complementary transcriptomic information, `scMultiNODE` recovers coherent developmental trajectories from scATAC-seq, a modality whose sparsity and weak cell-type separability often obscure trajectory structure when analyzed alone. `scMultiNODE` also identifies development-related, ATAC-specific genes and transfers annotations from RNA to unlabeled ATAC cells. Together, these results show that `scMultiNODE` produces an interpretable joint latent space that captures complex cell trajectories, supports the reconstruction of cell transition paths, and facilitates the discovery of development-related genes, which are key elements for studying cell differentiation. We envision that `scMultiNODE` will be critical to integrative analyses of multi-modal temporal single-cell datasets, especially those with unaligned measurements.

## 2. Methods

`scMultiNODE` aligns scRNA-seq (gene expression) and scATAC-seq (chromatin accessibility) datasets measured across time. We denote 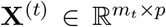 as the gene expression of *m*_*t*_ cells and *p* genes at timepoint *t* and 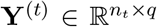 as the chromatin accessibility of *n*_*t*_ cells and *q* chromatin regions at timepoint *t*. Vectors are denoted in bold lowercase letters, and matrices are in bold capital letters. Given gene expression and chromatin accessibility matrices at observed timepoints *T*_RNA_, *T*_ATAC_ ⊆ {0, 1, 2, · · ·}, we aim to learn a unified low-dimensional representation that brings together multiple modalities, reflects cell type variations, and preserves dynamic cellular progression for downstream developmental analysis. We assume consistent feature sets across timepoints: the same genes for scRNA-seq and the same chromatin regions for scATAC-seq. While no prior cell-to-cell correspondence is needed, we assume the datasets share common biological signals, such as overlapping cell types and similar developmental trajectories, making joint integration both meaningful and feasible. `scMultiNODE` infers the necessary correspondence during learning and learns a single *d*-dimensional latent representation (*d* ≪ min{*p, q*}) that we require to: (1) place biologically similar cells from the two modalities close together, so that local cell-type structure is shared rather than confined to one modality, and (2) arrange cells so that their representations reflect continuous global progression over time.

### 2.1. Scalable integration while preserving local cell relationships

In this multi-modal integration problem, the important step is to align two single-cell modalities measured separately and representing distinct feature spaces. To achieve this, we use Gromov-Wasserstein (GW) optimal transport^27^ to align cells across modalities in an unsupervised manner.

GW optimal transport aims to move data points from one feature space to another while preserving the original local geometry captured by intra-modality distances. The central concept of GW is to find the best data correspondence matrix **T** that denotes the alignment between each data point across two data modalities, through minimizing the transport cost (i.e., the GW distance) as

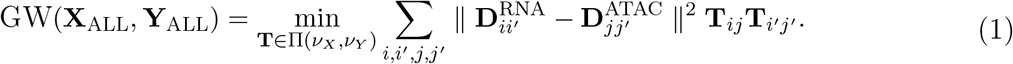

Here, **X**_ALL_ = CONCAT(**X**^(*t*)^|*t* ∈ *T*_RNA_) and **Y**_ALL_ = CONCAT(**Y**^(*t*)^|*t* ∈ *T*_ATAC_) are concatenations of gene expression and chromatin accessibility of all cells across available timepoints. *ν*_*X*_ and *ν*_*Y*_ are empirical distributions of **X**_ALL_ and **Y**_ALL_, respectively. **D**^RNA^ and **D**^ATAC^ are intra-modality distance matrices. Due to high dimensionality and sparsity of single-cell data, `scMultiNODE` employs an autoencoder (AE)^28^ to map each modality to its latent variable **Z** and construct intra-modality distance matrices in the latent space with k-nearest neighbor (kNN) distance:

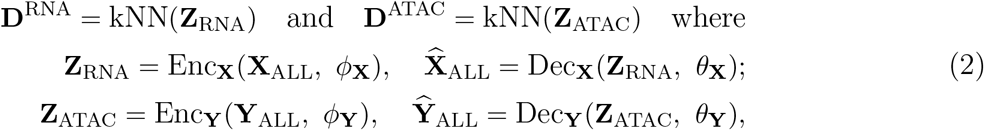

with encoder Enc(·, *ϕ*) and decoder Dec(·, *θ*) for each modality.

In the cell correspondence matrix **T**, the element **T**_*ij*_ indicates the alignment probability of RNA cell *i* and ATAC cell *j*. The GW loss function measures how intra-modality cell-pair distances in one dataset compare with those in the other dataset. So, the GW distance does not directly compare data points across different feature spaces, but instead finds a cross-space sample that best preserves intra-modality structure. This ensures that the transport plan **T** can preserve local geometry, like cell clusters and cell-cell neighboring relationships, in one modality and correspond to similar local structures in the other when we align two single-cell modalities. We binarize the predicted **T** such that for a RNA cell *i*, **T**_*ij*_ = 1 if ATAC cell *j* has the highest alignment probability with *i*; otherwise **T**_*ij*_ = 0. Notice that `scMultiNODE` does not require cell correspondence information as a prior and predicts it in an unsupervised manner^a^.

While previous unsupervised integration methods have successfully aligned single cells using GW optimal transport,^27^ the exact computation of the GW distance (Eq. 1) is NP-hard and incurs high computational costs for large-scale single-cell datasets. Therefore, `scMultiNODE` utilizes Quantized Gromov-Wasserstein (QGW),^26^ based on a divide-and-conquer strategy that partitions feature spaces into several blocks and aligns blocks recursively, to significantly speed up computations and scale to a large number of cells (Supp. Sec. S1).

With the help of the inferred cell correspondence **T**, `scMultiNODE` can now integrate gene expression and chromatin accessibility measurements into a joint latent space. Specifically, `scMultiNODE` uses a neural network Fus(·, *ω*) : ℝ^*d*^ i → ℝ^*d*^ parameterized by *ω*, named the fusion layer, to map modality-specific cell latent variables of each modality to the *d*-dimensional joint latent space through

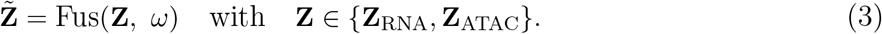

We hypothesize that if cells *i* and *j* from different modalities are biologically similar, they should be aligned in the joint latent space and their latent representations 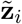 and 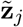 should be close to each other. To this end, `scMultiNODE` aligns modalities in the joint latent space by minimizing the loss function

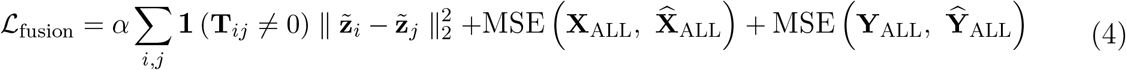

with the indication function **1**(·). Latent variables 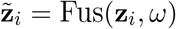 and 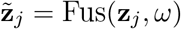 for RNA cell *i* and ATAC cell *j* (from Eq. 3), and

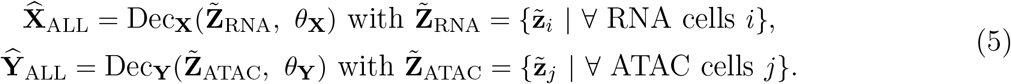

Therefore, during fusion layer training, we minimize the distance between the latent representations of cell pairs matched by the GW optimal transport alignment (i.e., **1** (**T**_*ij*_≠ 0)), weighted by the hyperparameter *α*. Since the decoders operate on the joint latent space, we also include the MSE reconstruction loss in the fusion objective (Eq. 4) to update the model.

### 2.2. Incorporating global cellular dynamics with neural ODE

`scMultiNODE` uses QGW to capture local cell relationships in the joint latent space. However, the joint space for now does not account for global cellular dynamics, such as developmental trajectories or lineage bifurcations. All the previous integration methods have this limitation, hence, their latent representations poorly define natural cell paths across timepoints and hinder downstream analysis.

To address the problem, `scMultiNODE` uses neural ODEs^29^ to explicitly model cell developmental dynamics in the joint latent space. Neural ODEs parameterize the temporal evolution of latent cell states by learning a neural network–based derivative function. While existing flow-based frameworks^9,30^ have used neural ODEs (with respect to time) to construct continuous-time trajectories and model single-cell development from gene expression data, `scMultiNODE` extends them to model the aligned representations of multi-modal datasets. It quantifies changes of cell latent representation 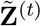 in the joint latent space at time *t* through a neural ODE: 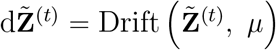. Here, 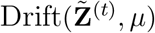 is a single non-linear neural network parameterized by *µ* and shared across both modalities, modeling the instantaneous rate of change of any cell’s latent state in the joint space. `scMultiNODE` calculates the initial condition 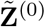 of cells at the first time *t* = 0, defined as the earliest observed timepoint for both modalities. `scMultiNODE` then predicts the subsequent cell states step-wise at any timepoint *t* through (assume RNA modality has the first timepoint as 0 ∈ *T*_RNA_)

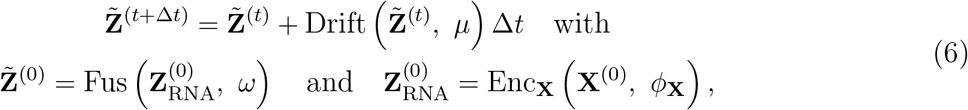

Here, hyperparameter Δ*t* denotes step size. We use the first-order Euler method (in Eq. 6) for convenience of explanation and one can specify any ODE solver in our implementations.

To fit the continuous trajectory (controlled by Drift neural network) to the observations, `scMultiNODE` minimizes the difference between the input and the reconstructed data. Specifically, at each measured timepoint *t* ∈ *T*_RNA_ or *t* ∈ *T*_ATAC_, `scMultiNODE` uses the decoder Dec_**X**_/Dec_**Y**_ to convert latent variables 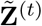 generated from Eq. 6 back to the high-dimensional feature space. Because we have no correspondence between true cells and cells generated from the ODE model, `scMultiNODE` utilizes the 2-Wasserstein metric^31^ to measure the distance between distributions defined by ground truth **X** and predictions 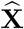 as

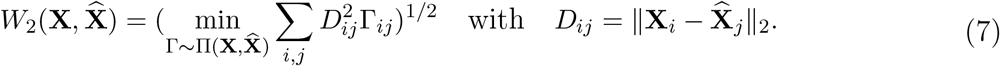

Wasserstein distance measures the distribution distance in the same metric space, different from the GW version that is applied for aligning different metric spaces. Here, 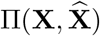 denotes the set of all transport plans between each cell of **X** and 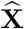 and *D*_*ij*_ represents the *ℓ*_2_ distance, such that the Wasserstein metric adopts the minimal-cost transport plan Γ to measure the data dissimilarity. `scMultiNODE` utilizes Wasserstein distance as the reconstruction loss when training the neural ODE.

Furthermore, to incorporate the cellular dynamics of neural ODE, `scMultiNODE` uses a dynamic regularization term to update the joint latent space with global developmental dynamics. The dynamic regularization is proposed in our previous work^9^ for modeling temporal scRNA-seq data, which incorporates cellular dynamics into the latent space to make it more robust to distribution shifts in the measurements across time. Here, we extend dynamic regularization in the multi-modal integration setting. Specifically, the dynamic regularization minimizes the difference between the joint latent representations generated by the fusion layer (i.e., Fus (Enc_**X**_(**X**, *ϕ*_**X**_)) or Fus (Enc_**Y**_(**Y**, *ϕ*_**Y**_))) and the dynamics learned by the ODE (i.e., 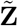). Because we have no correspondence between them, `scMultiNODE` again uses Wasserstein distance to evaluate their difference at each timepoint and defines the dynamic regularization:

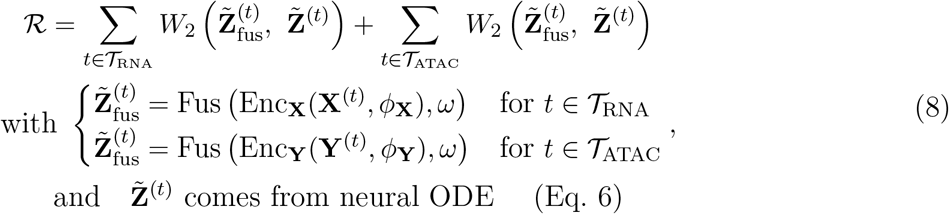

This term folds the learned dynamics back into the shared representation. Because the constraint applies identically to both modalities, a cell whose own features weakly determine its latent position is nonetheless drawn toward the shared, global developmental flow.

With these, `scMultiNODE` jointly optimizes all components by minimizing the regularized loss function

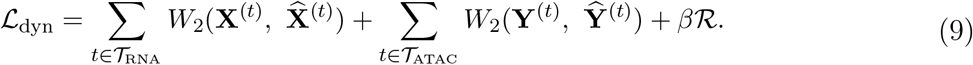

By embedding cellular dynamics, `scMultiNODE` improves upon previous integration models. This enables more effective data fitting and yields a joint latent space that is more robust and interpretable by capturing both global cellular dynamics and local cell relationships. Pseudocode for `scMultiNODE` is provided in Supp. Algorithm S1.

## 3. Experiment Setup

### Datasets

We use four developmental single-cell datasets with scRNA-seq and scATAC-seq measured at multiple timepoints across several species and tissues (Supp. Table S1): two coassay datasets — human cortex (HC)^12^ and human organoid (HO);^5^ two unaligned datasets — drosophila embryogenesis (DR),^13^ and mouse neocortex (MN).^32^ For each modality, we keep the top 2,000 most variable features and use batch-corrected data (Supp. Sec. S3).

### Baselines

We compare `scMultiNODE` with six state-of-the-art unsupervised single-cell integration methods that are capable of aligning different single-cell measurements (details in Supp. Sec. S4): `Seurat`,^19^ `UnionCom`,^33^ `uniPort`,^23^ `Pamona`, `SCOTv1`,^22^ and `SCOTv2`.^18^

### Evaluation metrics

We evaluate each model’s integrated latent representations from multiple perspectives (details in Supp. Sec. S5): *modality integration, cell type preservation, cell label transfer*, and *cellular dynamics*. Modality integration is assessed using `batch entropy`,^23^ fraction of samples closer than the true match (`FOSCTTM`),^22^ `neighborhood overlap`,^33^ and Spearman correlation coefficient (`SCC`). Cell type preservation is quantified by normalized mutual information (`NMI`)^34^ between Louvain clustering^35^ labels in the integrated latent space and ground truth cell type. Cell label transfer is evaluated by *k*NN-based label transfer accuracy using cell type labels (`LTA-type`).^18,23,33^ To assess cellular dynamics, we compute label transfer accuracy using timepoint labels (`LTA-time`) and the distance correlation between latent representations and timepoint labels (`time correlation`).

### Implementation

Hyperparameters for all methods are optimized using Optuna^36^ over broad search ranges (Supp. Sec. S6) and use latent dimension 50. `scMultiNODE`, we use a first-order Euler solver with step size Δ*t* = 0.1. We conduct ablation studies of `scMultiNODE` and give heuristic guidance on how to set hyperparameters in real-world scenarios in Supp. Sec. S9.

## 4. Experiment Results

### 4.1. scMultiNODE integrates modalities without sacrificing temporal or rare-population structure

#### Static integration discards temporal structure while scMultiNODE preserve it

The quality of the integrated latent space is first assessed using 2D Uniform Manifold Approximation and Projection (UMAP)^37^ and Principal Component Analysis (PCA) projections (Supp. Fig. S1-S4). The visualizations show that `scMultiNODE` effectively aligns the two modalities across all timepoints while preserving cellular dynamics. In contrast, baseline methods either fail to achieve effective cross-modal alignment over time (e.g., `Pamona` and `UnionCom`) or neglect temporal structure (e.g., `SCOTv2`), resulting in latent spaces dominated by cell type signals. Across all datasets, `scMultiNODE` consistently preserves both cell type variation and temporal cellular dynamics during integration.

Regarding the `batch entropy` score, `scMultiNODE` consistently outperforms the baselines across all datasets (Fig. 2A and E). Furthermore, `scMultiNODE` achieves the highest `time correlation` across all datasets (Fig. 2C and G), demonstrating that it aligns the two modalities well while preserving the underlying dynamics.

**Fig. 2.**
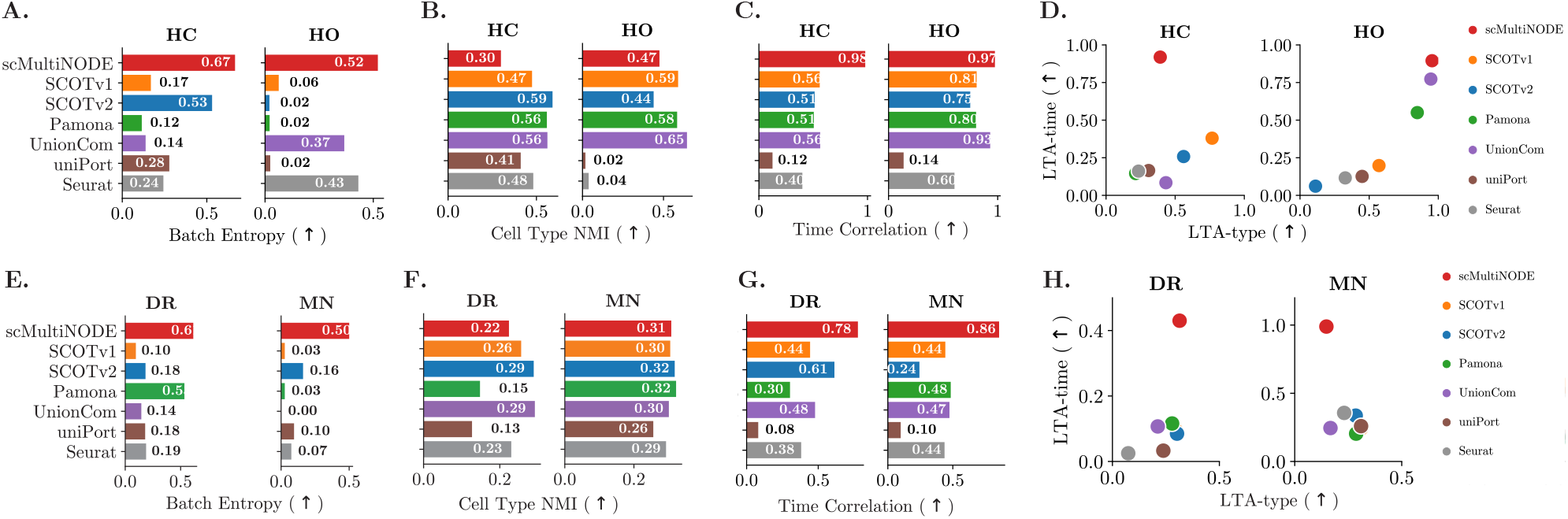
Integration evaluation on all datasets. (**A** and **E**) `Batch entropy` as an example of modality integration performance. (**B** and **F**) Cell type `NMI` score. (**C** and **G**) `Time correlation` evaluates integrations on capturing cell dynamics. (**D** and **H**) Label transfer accuracy (LTA) across modalities. We plot the `LTA-type` /`LTA-time` scores on the X/Y axes for all the methods, respectively.

Then, we validate whether the integration preserves distinct cell type separations (Fig. 2B and F). `scMultiNODE` achieves cell-type clustering performance comparable to existing baselines across most datasets. For instance, `scMultiNODE` attains `NMI` =0.47 on HO dataset (baseline median `NMI` =0.51) and `NMI` =0.31 on MN dataset (baseline median `NMI` =0.30). Plotting `LTA-type` against `LTA-time` (Fig. 2D and H) demonstrates that `scMultiNODE` uniquely occupies the upper region of both axes, achieving the highest `LTA-time` at a comparable `LTA-type` level. In contrast, baseline methods fail to simultaneously preserve temporal and cell-type heterogeneity. These results suggest that `scMultiNODE` sacrifices minimal static celltype separation to recover developmental trajectories that are overlooked by static integration methods. While incorporating cell-type supervision may improve clustering metrics, we find that it compromises modality alignment and temporal continuity (Supp. Sec. S8), likely due to rigid class constraints or noisy annotations that distort the latent space. Therefore, we adopt an unsupervised integration strategy throughout our experiments to better capture continuous cellular transitions. A complete list of metric values is provided in the Supplementary Note.

#### Single-modal dynamics are asymmetric across modalities while scMultiNODE overcome it

To assess the benefit of jointly modeling RNA and ATAC with `scMultiNODE`, we compared it against `scNODE`,^9^ a single-modal dynamics model trained separately on RNA (`scNODE (RNA)`) and ATAC (`scNODE (ATAC)`) across all datasets (Fig. 3A-B). Notably, `scNODE` performance is highly asymmetric, with `scNODE (ATAC)` substantially underperforming `scNODE (RNA)`, reflecting the sparser, noisier signal in ATAC data. In contrast, `scMultiNODE` largely closes this gap, significantly outperforming `scNODE (ATAC)` in `time correlation` while achieving the comparable or even better `time correlation` than `scNODE (RNA)`. A similar pattern holds for `NMI`, where `scMultiNODE` outperforms `scNODE (ATAC)` on three of four datasets, while remaining comparable to `scNODE (RNA)` on HO, the same tradeoff noted above. These results indicate that `scMultiNODE` ‘s key advantage lies in leveraging shared cross-modal structure to substantially improve dynamics and cell-type resolution for the weaker modality, without compromising performance on the stronger RNA modality.

**Fig. 3.**
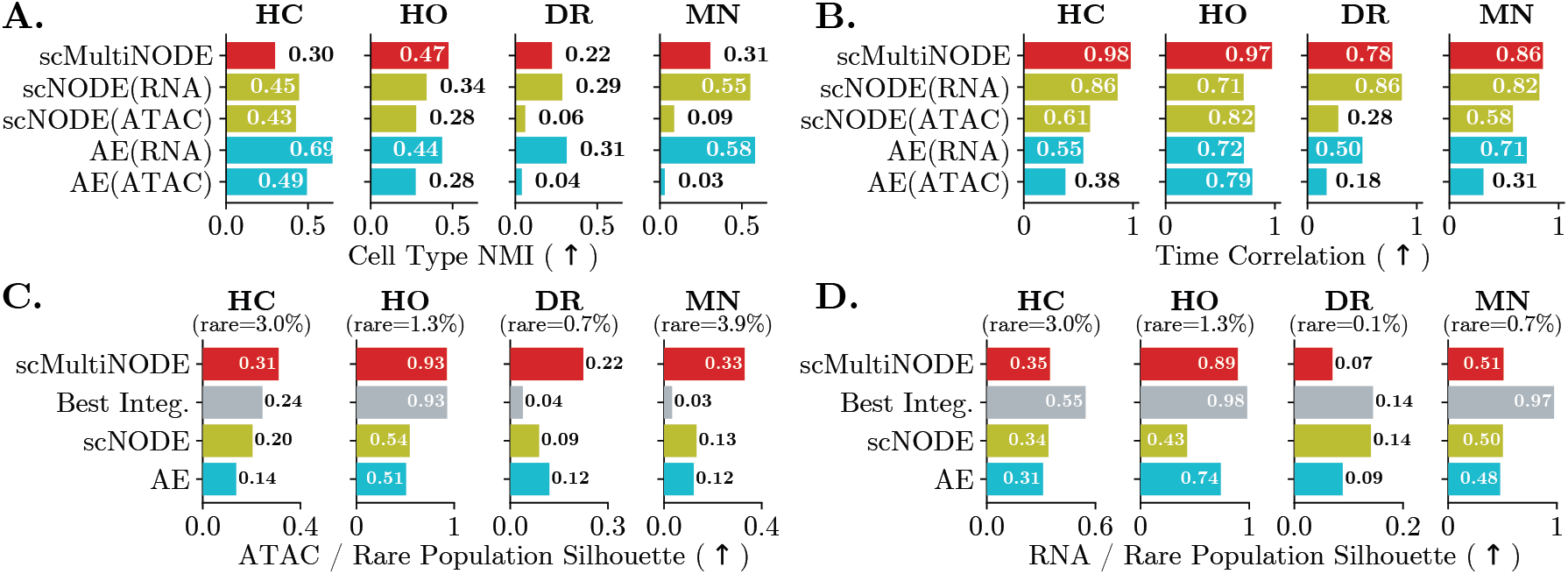
(**A, B**) We compare `scMultiNODE` with scNODE and AE trained on each modality separately in terms of cell-type `NMI` and `time correlation`. (**C, D**) Silhouette score of rare cell populations in the ATAC (**C**) and RNA (**D**) modalities, where `scMultiNODE` is compared against the best integration baseline, scNODE, and AE. The fraction of rare cells is annotated above each panel (“Best Integ”: best integration baseline model).

#### scMultiNODE rescues rare populations in the weak modality

Sparsity and low feature specificity make rare populations hard to resolve in scATAC-seq, limiting downstream analysis. To evaluate rare-population separation, we first perform integration and then compute a one-vs-rest silhouette score for the rare cell type (i.e., cell types with the fewest cells) in the learned joint latent space for ATAC and RNA (Fig. 3C-D).

On the ATAC modality, methods that incorporate either multi-modal integration or cellular dynamics outperform the single-modal AE, indicating that each component independently improves rare-population separation. The best integration baseline improves over the single-modal AE on some datasets (e.g., 0.24 of best integration vs. 0.14 of AE on HC), but collapses on others and fall below the AE (e.g., DR and MN), where the integration is dominated by the stronger RNA modality at the expense of ATAC. Modeling cellular dynamics alone (i.e., `scNODE`) yields only modest gains (e.g., 0.54 vs. 0.51 on HO), suggesting that neither integration nor dynamics alone is sufficient to overcome the challenges of weak modalities. By jointly modeling cross-modal correspondence and cellular dynamics, `scMultiNODE` achieves the highest silhouette score on all datasets. For example, `scMultiNODE` attains a silhouette score of 0.31 on HC, compared with 0.24 for the best integration baseline and 0.20 for scNODE (Fig. 3C). The improvement is even more significant on unaligned MN data, where `scMultiNODE` achieves 0.33 while all other methods remain below 0.15. Notably, `scMultiNODE` is the only method that avoids ATAC collapse on these datasets, highlighting the importance of jointly modeling integration and dynamics for reliable rare-population separation in weak modalities.

Although `scMultiNODE` yields the largest improvements on the weaker ATAC modality, it remains competitive on RNA (Fig. 3D). Relative to the matched single-modal RNA representation, `scMultiNODE` retains or improves rare-population separability. It is only surpassed by the dedicated integration baselines on RNA, but those same methods collapse on ATAC. This reflects a expected trade-off that because RNA already provides strong cell-identity signals, these baselines can achieve high RNA performance by effectively discarding ATAC, whereas `scMultiNODE` learns a balanced representation that resolves rare populations across both modalities. These results demonstrate that combining multi-modal integration with dynamic modeling enables `scMultiNODE` to more reliably resolve rare cell populations, particularly in weaker modalities such as scATAC-seq.

Together, these results demonstrate the key advantages of `scMultiNODE`. It preserves cell-type structure while better capturing temporal dynamics than static integration methods, reduces the RNA-ATAC performance gap observed in single-modal models, and more effectively resolves rare cell populations in the weak modality. Ablation studies further show that `scMultiNODE` is robust to latent space size but critically depends on dynamic regularization, whose removal breaks down dynamic modeling. Despite this added complexity, `scMultiNODE` remains computationally efficient and scalable (Supp. Sec. S9).

### 4.2. Joint dynamics recover multifurcating trajectories that neither modality supports alone

We demonstrate that `scMultiNODE` learns an interpretable joint latent space that preserves complex cellular developmental trajectories in multi-modal data. Previous methods typically infer pseudotime from the integrated latent space to reconstruct cell lineages.^19^ However, because they rely on static latent representations without explicitly modeling cellular dynamics, they often struggle to capture complex cell trajectories.

We conduct the evaluation on the human cortex (HC) dataset, which contains multifurcating developmental trajectories. Pseudotime is inferred from each latent space using Monocle3^38^ and evaluated by the Spearman correlation (*ρ*) with the ground-truth trajectory progression (Fig. 4A). We compare `scMultiNODE` with multi-modal integration baselines lacking dynamic modeling, as well as single-modality AE and scNODE models trained separately on RNA and ATAC. ATAC provides a particularly challenging test due to its sparsity and weak cell-type separation, allowing us to assess whether integration improves trajectory recovery in a weak modality.

**Fig. 4.**
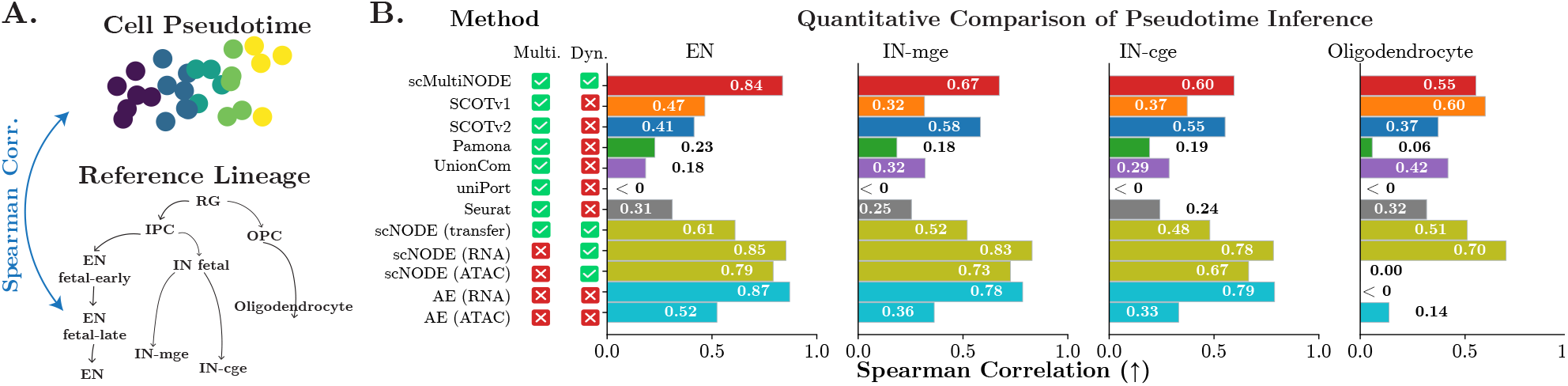
(**A**) We evaluate the consistency between cell pseudotime and reference cell lineage on the HC dataset. The trajectory originates from RG and branches into four lineages. (RG: Radial Glia, IPC: Intermediate Progenitor Cells, OPC: Oligodendrocyte Precursor Cells, EN: Excitatory Neurons, IN: Inhibitory Neurons) (**B**) Spearman correlation coefficient (*ρ*) between ground truth cell state orders and Monocle3 pseudotime of the integration “Multi.” indicates multi-modal integration method, “Dyn.” denotes models incorporating cell dynamics).

To test whether `scMultiNODE` ‘s performance can be achieved with a simpler two-step pipeline, we also evaluate `scNODE (transfer)`. Here, `scNODE` is trained on one modality to infer pseudotime, which is then transferred to the other modality via regression after alignment with each integration baseline (Supp. Sec. S11). We evaluate both RNA-to-ATAC and ATAC-to-RNA transfer, reporting the best result across all integration baselines.

Methods that model dynamics explicitly (`scMultiNODE` and `scNODE` (RNA)) preserve global trajectory structure, whereas static integration baselines produce collapsed or misaligned branches. For example, `scMultiNODE` reaches *ρ* = 0.84 on the EN lineage against 0.47 for the best integration baseline. On the Oligodendrocyte lineage `scMultiNODE` has *ρ* = 0.55, comparable to SCOTv1 (*ρ* = 0.60) and above all others (Fig. 4B).

The uplift on ATAC is the sharpest result. From ATAC alone, no baseline recovers the branching structure. The static AE reaches *ρ* = 0.14 (Oligodendrocyte) and 0.52 (EN), and `scNODE` (ATAC) fails on the Oligodendrocyte lineage. By contrast, `scMultiNODE` attains *ρ* = 0.55 (Oligodendrocyte) and *ρ* = 0.84 (EN), recovering coherent trajectories from a modality that does not support accurate inference on its own. The two-step pipeline is not sufficient. `scNODE (transfer)` improves on static integration (*ρ* = 0.61 vs the best integration baseline’s 0.47 on EN) by carrying learned dynamics across the alignment, but underperforms `scMultiNODE` on every branch and falls below single-modal `scNODE` in several cases, because errors in the intermediate alignment propagate into the transferred pseudotime. Dynamics and integration must be learned jointly rather than composed. We additionally perform this analysis using PAGA,^39^ a topology-based pseudotime inference method, and obtain similar results, indicating that `scMultiNODE` integration preserves not only trajectory direction but also complex topological structures (Supp. Fig. S7).

These results demonstrate that explicitly modeling global cellular dynamics and jointly integrating modalities are essential for accurate trajectory inference. Rather than simply adding RNA information, `scMultiNODE` leverages its joint architecture to transfer developmental dynamics through a shared neural ODE, enabling weak modalities such as ATAC to recover complex trajectories that cannot be resolved independently.

Applying Least Action Path (LAP) analysis^30^ on the `scMultiNODE` integration, we extracted coherent trajectories toward oligodendrocytes and glutamatergic neurons (Supp. Fig. S11). Genes identified along these trajectories show stronger temporal variation than random controls and include known developmental regulators. Notably, *SV2B* is absent from RNA-derived marker lists and appears only as an ATAC-specific marker, such that weak modality contributes signal that the strong one cannot see, highlighting the complementary biological information recovered through multi-modal integration (details in Supp. Sec. S12).

### 4.3. Label transfer makes unannotated chromatin data interpretable

Germ layers are foundational embryonic structures that give rise to diverse organs and cell types, making their identification essential for understanding development.^40^ While scRNA-seq provides strong transcriptional signals for annotation, scATAC-seq remains challenging due to limited labels.^11^ This unaligned setting complicates existing approaches for modeling biological processes across modalities. To address this challenge, we leverage the integration learned by `scMultiNODE` to transfer germ layer annotations from RNA to ATAC, enabling interpretation of the weakly annotated ATAC modality (Supp. Sec. S13).

We evaluated label transfer on the unaligned DR dataset, where germ layer labels were available only for RNA. A random forest classifier trained on RNA embeddings accurately predicted ATAC labels in `scMultiNODE`’s joint latent space, producing coherent germ layer clusters consistent with developmental trajectories (Supp. Fig. S12). Marker gene analysis of ATAC-derived gene activity confirmed clear germ layer separation (Supp. Fig. S13), whereas baseline methods showed poor resolution. Gene Ontology (GO) enrichment further validated biological relevance, revealing germ layer–specific functions (Supp. Fig. S14), such as nervous system development in the ectoderm group.^41^

Overall, `scMultiNODE` preserves developmental structure across modalities and enables accurate cross-modal label transfer, enhancing the interpretability of chromatin accessibility data and supporting developmental analyses when direct labels are unavailable.

## 5. Conclusion

`scMultiNODE` is a novel framework for integrating temporally resolved, multi-modal single-cell datasets across unaligned modalities and timepoints. It learns a latent space that preserves cell identity and developmental dynamics, improving trajectory inference and cross-modal label transfer. `scMultiNODE` outperforms existing methods while remaining computationally efficient and readily extensible to additional modalities.

## Acknowledgments

This project was funded by the National Institutes of Health (NIH) award 1R35HG011939-01.

## Footnotes

1 If cell correspondence is known (even partially) ahead, we can encode such information into **T** and skip the alignment procedure. Our experiments assume no cell correspondence is given.

