## Supplementary Notes for "scMultiNODE: Recovering Developmental Dynamics in Weakly Resolved Modalities through Multi-Modal Integration"

### S1 Scalable Gromov-Wasserstein Alignment

`scMultiNODE` uses Gromov-Wasserstein (GW) to align cells from different modalities. The exact computation of GW distance is NP-hard and requires expensive computational costs for large-scale single-cell datasets. Therefore, `scMultiNODE` utilizes a recently proposed approximation algorithm, Quantized Gromov-Wasserstein (QGW) [8], to significantly speed up computations. QGW uses a divide-and-conquer strategy, where it partitions feature spaces into several blocks and matches blocks recursively. We derive the QGW computation following Eq. 1 in the manuscript.

Specifically, QGW assumes the feature space $\mathbf{Z}$ can be divided into disjoint and nonempty sets $U_1, \cdots, U_r$, with $r << |\mathbf{Z}|$. For example, the large-scale single-cell measurements can be divided into several cell-type groups. Within each partition $U_s$ $(s = 1, \cdots, r)$, there exists a representative point $z_s \in U_s$, like the anchor cell for each cell type. We define $Z_r := \{z_1, \cdots, z_r\}$ as quantized representations of space $\mathbf{Z}$. Therefore, given latent representations $\mathbf{Z}_{\text{RNA}}$ and $\mathbf{Z}_{\text{ATAC}}$ and their quantized representations $Z_r^{\text{RNA}}$ and $Z_r^{\text{ATAC}}$, QGW algorithm proceeds in three steps.

- *Global Alignment:* QGW first computes a transport plan $\pi_r \in \mathbb{R}^{r \times r}$ between representative points of two modalities. Since the number of partitions $r$ is chosen to be much smaller than the number of cells, computing $\pi_r$ can be feasibly approximated via any existing GW algorithm. This transport plan provides a global alignment between partitioned blocks.
- *Local Alignment:* Then QGW produces a collection of local alignments between individual cells. For each partition $z_a \in Z_r^{\text{RNA}}$ and $z_b \in Z_r^{\text{RNA}}$, we compute the transport plan $\pi_{ab} \in \mathbb{R}^{|U_a| \times |V_b|}$ between the RNA block $U_a$ and ATAC block $V_b$ by solving

  $$\min_{\pi_{ab} \in \Pi(\nu_{U_a}, \nu_{V_b})} \sum_{i \in U_a, j \in V_b} \| \mathbf{D}_{iz_a}^{\text{RNA}} - \mathbf{D}_{jz_b}^{\text{ATAC}} \|^2 \pi_{ab}(i,j), \tag{S1}$$

  where $\mathbf{D}_{iz_a}^{\text{RNA}}$ is the distance between any cell of block $U_a$ and its representative point $z_a$ ($\mathbf{D}_{jz_b}^{\text{ATAC}}$ is defined in the same way). $\pi_{ab}(i,j)$ is the element at row $i$ and column $j$. [8] has proved that Eq. S1 can be solved efficiently with a log-linear time complexity with respect to the size of the maximum block.
- *Coupling:* With local and global transport plans, we can merge them into the final solution of QGW as

  $$\mathbf{T} = \sum_{a,b} \pi_r(a,b) \pi_{ab}. \tag{S1}$$

  [8] has proved that the QGW can give a good approximation when the feature spaces being compared admit compact partitions, which is a realistic setting for single-cell measurements where we have multiple homogeneous cell-type groups.

An important routine in QGW is generating good partitions. Since our `scMultiNODE` is an unsupervised model without cell type labels, we followed QGW defaults [8] as randomly and uniformly choosing a small set of cells without replacement as representative points and computed a Voronoi partition.

### S2 `scMultiNODE` Training Details

Our `scMultiNODE` is implemented with *Pytorch 1.13* [22] and is trained end-to-end. `scMultiNODE` training consists of three main steps. `scMultiNODE` first trains the AE components for each modality with all cells. We use Adam optimizer to train AEs by minimizing RNA/ATAC reconstruction MSE loss with a learning rate of 0.001 and 1000 iterations. Then, `scMultiNODE` aligns modality-specific latent representations with QGW optimal transport. In the QGW algorithm, we construct the intra-modality distance matrices $\mathbf{D}^{\text{RNA}}$

and $\mathbf{D}^{\text{ATAC}}$ through the kNN graph. Once the cell correspondence matrix $\mathbf{T}$ is estimated from the QGW algorithm, `scMultiNODE` maps modality-specific latent representations to a joint latent space (through fusion layer $\text{Fus}(\cdot, \omega)$) by minimizing $\mathcal{L}_{\text{fusion}}$ through the Adam optimizer with a learning rate of 0.001 and 1000 iterations. Finally, `scMultiNODE` adopts neural ODE to model the cellular dynamics and incorporate the learned dynamic into the joint latent space by minimizing $\mathcal{L}_{\text{dyn}}$. We adopt batch training and use the Adam optimizer to train `scMultiNODE` with a learning rate of 0.001 and 2000 iterations. At each training iteration, we randomly select 64 cells at $t = 0$ as a batch and predict for every timepoint $t \in \mathcal{T}_{\text{RNA}} \cup \mathcal{T}_{\text{ATAC}}$. Because the Wasserstein distance computation is expensive, batch training improves training efficiency and enables `scMultiNODE` usage on large-scale datasets. We use *geomloss* [12] to compute Wasserstein distance with blur = 0.05 and scaling = 0.5. Since the datasets we used contain the first time point for both modalities, we use the RNA modality at the first time point to compute the initial condition of the neural ODE. In our implementation, users can specify which modality at the first time point is used to initialize the neural ODE.

**Algorithm S1** `scMultiNODE`

**Input:** The set of timepoint indices $\mathcal{T}_{\text{RNA}}$ and $\mathcal{T}_{\text{ATAC}}$; gene expression matrices $\{\mathbf{X}^{(t)} \mid t \in \mathcal{T}_{\text{RNA}}\}$; chromatin peak/gene activity matrices $\{\mathbf{Y}^{(t)} \mid t \in \mathcal{T}_{\text{ATAC}}\}$; hyperparameters $\Delta t$, $\alpha$, $\beta$, the number of neighbors $k$ in kNN; randomly initialized neural networks $\text{Enc}_{\mathbf{X}}$, $\text{Enc}_{\mathbf{Y}}$, $\text{Dec}_{\mathbf{X}}$, $\text{Dec}_{\mathbf{Y}}$, fusion layer Fus, and the neural ODE drift network Drift.

**( Step I: dimensionality reduction )**
$\mathbf{X}_{\text{ALL}} = \text{CONCAT}\left(\mathbf{X}^{(t)} \mid t \in \mathcal{T}_{\text{RNA}}\right)$
$\mathbf{Y}_{\text{ALL}} = \text{CONCAT}\left(\mathbf{Y}^{(t)} \mid t \in \mathcal{T}_{\text{ATAC}}\right)$
Optimize RNA-related AE parameters $\phi_{\mathbf{X}}$ and $\theta_{\mathbf{X}}$ to minimize MSE for RNA
Optimize ATAC-related AE parameters $\phi_{\mathbf{Y}}$ and $\theta_{\mathbf{Y}}$ to minimize MSE for ATAC

**( Step II: modality integration )**
Construct intra-modality distance matrices $\mathbf{D}^{\text{RNA}}$ and $\mathbf{D}^{\text{ATAC}}$ with representations $\mathbf{Z}_{\text{RNA}}$ and $\mathbf{Z}_{\text{ATAC}}$
Use Quantized Gromov-Wasserstein (QGW) algorithm to predict cell correspondence matrix $\mathbf{T}$
Map latent representations to joint latent space through fusion layer
Optimize fusion layer parameter ($\omega$) and AE parameters to minimize $\mathcal{L}_{\text{fusion}}$

**( Step III: cellular dynamics )**
Optimize the entire model to minimize $\mathcal{L}_{\text{dyn}}$

**Output:** modality integration $\tilde{\mathbf{Z}}_{\text{RNA}} = \text{Fus}(\mathbf{Z}_{\text{RNA}}, \omega)$ and $\tilde{\mathbf{Z}}_{\text{ATAC}} = \text{Fus}(\mathbf{Z}_{\text{ATAC}}, \omega)$

### S3 Single-Cell Dataset and Pre-Processing

We use four publicly available developmental single-cell datasets with scRNA-seq and scATAC-seq assays to demonstrate the capabilities of `scMultiNODE` in integrating modalities in an unsupervised manner.

- **Human cortex (HC):** Zhu et al. generate transcriptomic and chromatin accessibility data using multi-omic single-nucleus RNA sequencing (snRNA-seq) and single-nucleus assay for transposase-accessible chromatin (snATAC-seq). The dataset profiles 45549 cells in total across a broad developmental time frame from human fetal cortical plate to adult specimens [36]. They have normalized data with scTransform [14] and removed batch effects. We use the processed data provided in its original paper, which contains normalized gene expression count data, and the gene activity matrix inferred from ATAC-seq that assesses chromatin accessibility at the gene body and promoter regions. We randomly sample 5% of cells and test our model on this subset with 2277 cells. For this and the following datasets, we subsample cells while retaining cell type proportions. For each modality, we select the top 2000 highly variable genes (HVGs) using Scanpy [30]. The HC data can be downloaded from the CELLxGENE portal (https://cellxgene.cziscience.com/collections/ceb895f4-ff9f-403a-b7c3-187a9657ac2c).

- **Human organoid (HO):** Fleck et al. have acquired paired single-cell transcriptome (scRNA-seq) and accessible chromatin (scATAC-seq) data with 34088 cells over a dense time course (spanning 4 days to 2 months) of human brain organoid development [13]. The dataset collects brain organoids of the same batch that dissociated at multiple timepoints during brain organoid development. The original paper provides gene expression count data of RNA-seq and the gene activity matrix inferred from ATAC-seq. We randomly sample 10000 cells and test our model on this subset. For each modality, we also select the top 2000 HVGs. The HO data can be downloaded from Zenodo (https://zenodo.org/records/5242913).
- **Drosophila embryogenesis (DR):** Calderon et al. profile chromatin accessibility in almost 1 million nuclei and gene expression in half a million nuclei from eleven overlapping windows spanning the entirety of drosophila embryogenesis (0 to 20 hours) [3]. The dataset contains the scRNA-seq profile of 547805 cells and scATAC-seq measurements for 976460 cells. For each modality, we randomly sample 5% of cells and test our model on the gene expression count matrix of 2738 cells and chromatin peak matrix of 4246 cells. As in the Seurat workflow [16], we select the top 2000 HVGs of scRNA-seq data and the top 2000 variable peaks for scATAC-seq data. The original paper shows that the data are not confounded by batch effects. The DR data is downloaded from https://shendure-web.gs.washington.edu/content/members/DEAP_website/public/.
- **Mouse neocortex (MN):** Yuan et al. provide a single-cell dataset of transcriptional (scRNA-seq) and epigenomic (scATAC-seq) measurements over a time course spanning for mammalian neocortical neurons in both mouse and marmoset [34]. The batch effects across timepoints and mammalian libraries have been corrected with the Seurat package. We use the mouse neocortex data and randomly select 10% cells for both modalities, obtaining a gene expression count matrix of 6098 cells and a chromatin peak matrix for 1914 cells. We select the top 2000 HVGs of scRNA-seq data and the top 2000 variable peaks for scATAC-seq data using Scanpy and EpiScanpy [9]. The MN data is downloaded from Gene Expression Omnibus SuperSeries GSE204851 (https://www.ncbi.nlm.nih.gov/geo/query/acc.cgi?acc=GSE204851).

To make computations tractable, we relabel timepoints with consecutive natural numbers starting from 0. We normalize the gene expression count matrix to remove cell-specific bias before conducting experiments. Specifically, given the count expression of cell $i$ as $\mathbf{X}_i \in \mathbb{R}^p$, we normalize it by total counts over all genes

$$\mathbf{X}_i = \frac{\mathbf{X}_i}{\sum_{j=1}^{p} \mathbf{X}_i} * 10^4, \qquad \text{followed by} \qquad \mathbf{X}_{ij} = \log(\mathbf{X}_{ij} + 1). \tag{S3}$$

Because the HC dataset already provides normalized gene expression data, we normalize the scRNA-seq data matrix of the other datasets. Furthermore, we apply a standard preprocessing step to the chromatin accessibility matrix by binarizing the count values, such that each entry reflects the presence (1) or absence (0) of accessibility at a given peak in each cell. This approach helps mitigate technical noise inherent in the highly sparse chromatin accessibility data. An overview of these datasets is included in Supp. Table S1.

### S4 Baseline Models

We compare `scMultiNODE` with six state-of-the-art unsupervised single-cell integration methods that are capable of aligning multiple modalities and computing the joint latent space.

- `Seurat`: The single-cell analysis platform `Seurat` [16] projects two datasets into a common space with linear canonical correlation analysis (CCA) that maximizes cross-dataset correlation. `Seurat` first identifies correspondence anchor points via CCA and then imputes one modality to another modality based on anchors. We use Seurat v5 in our experiments.
- `SCOTv1`: Demetci et al. [11] present the unsupervised learning method SCOT to align single-cell multimodal datasets with Gromov-Wasserstein (GW) optimal transport. We term this model as `SCOTv1` in this paper. We use the `SCOTv1` implementation on https://github.com/rsinghlab/SCOT.
- `SCOTv2`: The `SCOTv2` [10] model improves upon `SCOTv1` by using unbalanced GW optimal transport to deal with disproportionate cell type representation and differing numbers of cells across single-cell modalities. We use the `SCOTv2` implementation on https://github.com/rsinghlab/SCOT.

- `UnionCom`: Cao et al. [5] propose `UnionCom`, another unsupervised multi-modal integration model. It matches two datasets based on geometrical matrix matching. Specifically, `UnionCom` computes intra-modality distance matrices and then matches the modalities based on a matrix optimization problem. We use the `UnionCom` implementation on https://github.com/caokai1073/UnionCom. Apart from the hyperparameters listed in the Supp. Table S2, we set all its other hyperparameters as default.
- `Pamona`: The `Pamona` [7] method adopts partial GW optimal transport to integrate multi-modal single-cell datasets. It aims to obtain shared and dataset-specific cell variations across modalities. We use the `Pamona` implementation on https://github.com/caokai1073/Pamona. We set the number of shared cells between datasets as the minimal number of cells across all modalities.
- `uniPort`: Cao et al. [6] introduce `uniPort`, incorporating coupled variational auto-encoders and mini-batch unbalanced optimal transport to integrate multi-modal single-cell datasets. We use its implementation on https://github.com/caokai1073/uniPort in our experiments. We use the diagonal integration mode for `uniPort`.

Furthermore, to assess the benefit of jointly modeling RNA and ATAC with `scMultiNODE` , we compare it against two single-modal baselines:

- `scNODE`: Zhang et al. [35] propose `scNODE`, a generative model that explicitly captures cellular dynamics in temporal scRNA-seq data. `scNODE` combines a variational autoencoder with a neural ODE to learn a latent space that models continuous developmental trajectories over time. Because it is a single-modal method, we train it independently on RNA and ATAC, yielding `scNODE(RNA)` and `scNODE(ATAC)`. We use the `scNODE` implementation on https://github.com/rsinghlab/scNODE.
- `AE`: As the simplest single-modal baseline, we train a standard autoencoder that embeds each modality into a low-dimensional latent space by minimizing the reconstruction MSE loss, without any cross-modal alignment or dynamic modeling. We train it independently on RNA and on ATAC, obtaining `AE(RNA)` and `AE(ATAC)`. They provide reference baselines for disentangling the contributions of modality integration and dynamics modeling.

### S5 Evaluation metrics

We evaluate each model's integration from four perspectives: modality integration, cross-modal cell label transfer, capturing cell type variation, and preserving cellular dynamics. Therefore, we adopt the following evaluation metrics.

**Modality integration.** We use `batch entropy` to evaluate the integration of all datasets. `Batch entropy` is originally introduced in Xiong et al. [32] and previously adopted by Cao et al. [6]. It evaluates the sum of regional mixing entropies between different datasets where a high score indicates cells from different modalities are mixed well. Specifically,

$$\text{batch entropy} = \sum_{k \in \{\text{RNA},\text{ATAC}\}} -p'_k \log\left(p'_k\right) \qquad \text{with} \qquad p'_k = \frac{p_i/P_i}{\sum_{j\in\{\text{RNA},\text{ATAC}\}} p_j/P_j}, \tag{S5}$$

in which $P_k$ is the proportion of cells in each modality, and $p_k$ is the proportion of cells from modality $k$ in a given region.

Specifically, for the co-assay datasets (HC and HO) where one-to-one cell correspondence information is implicitly available, we additionally use the fraction of samples closer than the true match (`FOSCTTM`) [20,11], `neighborhood overlap` [5], and Spearman correlation coefficient (`SCC`). For each data point in the joint latent space, `FOSCTTM` computes the fraction of data points that are closer than its true nearest neighbor (i.e., the matched cell). We average these fraction values for all the cells in both modalities. A perfect integration implies that all cells should be closest to their true match, resulting in a `FOSCTTM` of zero. Therefore, a lower `FOSCTTM` value denotes a better integration performance. Furthermore, the `neighborhood overlap` is defined similarly and computes the ratio of cells that can find their correspondence cells from the other dataset in their neighborhood. We use the averaged ratio of `neighborhood overlap` of the two modalities.

The `neighborhood overlap` ranges from 0 to 1, and a higher value implies a better recovery of cell-to-cell correspondence between the two modalities. Lastly, based on the intuitive assumption that matched cells should have similar latent representations in the joint latent space, we use `SCC` to evaluate representation similarities between matched cells, such that a better integration leads to a higher `SCC` value. In our analysis, we show `1-FOSCTTM` instead of `FOSCTTM` to unify metrics comparison, where a higher metric value implies better integration performance.

**Cell label transfer.** We also evaluate integration using cell type labels through label transfer accuracy (`LTA-type`) as in previous studies [5,6,10]. This metric assesses the clustering of cell types after integration by training a $k$-nearest neighbor (kNN) classifier on joint latent representations of one modality and then evaluates its predictive accuracy on another modality. It ranges from 0 to 1, and a higher metric value indicates better integration performance as cells that belong to the same cell type are aligned close together.

**Cell type variation.** To assess how well cell type clusters are preserved after integration, we calculate the Normalized Mutual Information (`NMI`) score [21]. It quantifies the similarity between predicted clusters and ground truth labels, ranging from 0 to 1, with higher values indicating better preservation of cell groups. Specifically, we apply the Louvain algorithm [2] to cluster cells in the integrated latent space and calculate the NMI between the resulting cluster labels and the true cell types.

**Cellular dynamics.** As the main objective of our research, we evaluate how well the integration captures the cellular variations across different timepoints. Therefore, we compute label transfer accuracy using timepoint labels (named as `LTA-time`), such that a higher `LTA-time` indicates better integration performance as cells that belong to the same timepoint are aligned close together. Furthermore, we hypothesize that good latent representations, if they retain the developmental dynamics, should highly correlate with the timepoint label. Therefore, we define the `time correlation`, which computes the distance correlation [28] between cell representations in the joint latent space and their corresponding timepoint labels. The distance correlation measures linear and nonlinear association between two datasets of arbitrary dimensions. Hence, the `time correlation` ranges from 0 to 1, where a higher value implies a better integration, which is highly associated with cellular dynamics. We use the `dcor` [27] package to compute distance correlations.

### S6 Hyperparameter Tuning

On co-assay datasets (HC and HO), we select corresponding hyperparameters for all methods (`scMultiNODE` and baselines) that yield the minimum `FOSCTTM` value; on unaligned datasets (DR and MN), we select hyperparameters that yield the maximum `LTA-type` based on common cell type labels. We use Optuna [1] to automatically determine the optimal hyperparameters and use sufficiently large hyperparameter ranges for search and evaluation. The hyperparameter search ranges of `scMultiNODE` and baselines are listed in Supp. Table S2. We set the joint latent space dimension as 50 for all methods. We use the first-order Euler ODE solver and set ODE step size $\Delta t = 0.1$ in `scMultiNODE` . We run each method for sufficient iterations to ensure they converge.

### S7 Comparison of Multi-Modal Integration

We compare `scMultiNODE` 's integration performances with baseline methods on both co-assay and unaligned datasets. The UMAP and principal component analysis (PCA) visualizations of integrations for all datasets are shown in Supp. Fig. S1-S4. All evaluation metric calculations were implemented in Python and the detailed metric values are listed in Supp. Fig. S5 and Table S3-S4.

### S8 `scMultiNODE` Training with Cell Type Supervision

To investigate the effect of incorporating cell type supervision into our `scMultiNODE` framework, we extend it by adding a classification head on top of the latent space. Specifically, we use a fully connected layer with a softmax activation to predict cell type labels, and train the classification head with cross-entropy loss. This cross-entropy loss is jointly optimized with the original loss function of `scMultiNODE` , thereby encouraging the latent representations to reflect known cell type groups. The classification loss is weighted by a tunable coefficient, set to 1.0 in this experiment. All other hyperparameters follow the tuning procedure described in Sec. S6. We apply this supervised variant of `scMultiNODE` to all datasets and compare to the unsupervised (original) version of `scMultiNODE` .

UMAP visualizations (Supp. Fig. S10) reveal that incorporating cell type supervision significantly improves the separation of cell groups in the joint latent space. For example, on the HC dataset, the supervised model achieves a `NMI` of 0.614, compared to 0.299 for the unsupervised version. The `LTA-type` also increases from 0.392 to 0.690, suggesting that cell type boundaries are more distinct when supervision is applied (Supp. Table S10-S11). However, this improvement in cell type clustering comes at the cost of modality integration and cellular dynamics preservation. The `batch entropy`, which measures how well cells from different modalities are integrated, decreases from 0.667 (unsupervised) to 0.453 (supervised) on HC dataset. Similarly, the `time correlation` drops from 0.979 to 0.911, and `LTA-time` declines significantly from 0.919 to 0.515 on HC data (Supp. Table S10-S11). These show that the supervised model captures less of the underlying developmental dynamics.

These results highlight a key trade-off. While cell type supervision improves clustering performance and facilitates cell group separation, it constrains the latent space in a way that limits cross-modality integration and dynamic structure preservation. This suggests that the supervised classification loss term may distort the geometry of the latent space by forcing it to align the discrete cell group boundaries, which may not reflect the continuous nature of biological processes. Additionally, inaccurate or coarse-grained cell annotations could introduce label noise, further biasing the learned representations. In our study here, we focus on unsupervised modality integration.

### S9 Investigation of `scMultiNODE` Hyperparameters

We evaluate `scMultiNODE` 's performance under different hyperparameter settings using two representative datasets: a co-assay dataset (HC) and an unaligned dataset (DR). We first run `scMultiNODE` with the joint latent space size $d$ varying from $\{10, 50, 100, 150, 200\}$. Supp. Table S6 shows that `scMultiNODE` is robust in terms of the size of the latent dimensionality. Users can choose to set a reasonable latent dimension based on a tradeoff between accuracy and computational costs. State-of-the-art methods [17] generally choose a latent space of 10 to 50 dimensions. For a fair comparison, we set the latent size $d = 50$ for all methods in our experiments.

`scMultiNODE` uses QGW optimal transport to align cell representations from two modalities and ensures aligned cells have similar latent representations. The QGW algorithm calculates the intra-modality distance matrix using kNN. Thus, we vary the number of neighbors $k$ to be considered in kNN from $\{5, 10, 50, 100, 150, 200\}$. As shown in Supp. Table S7, `scMultiNODE` outperforms the best baseline model in terms of integration quality, with little impact from changing the number of neighbors. Additionally, the coefficient $\alpha$ for the matched cell integration loss (in Eq. 4) is varied from $\{0.0, 0.01, 0.1, 1.0, 10.0, 100.0\}$. Supp. Table S8 indicates that performance drops noticeably when $\alpha = 0$, where matched cells are not encouraged to converge in the latent space. Specifically, on the DR dataset, `scMultiNODE` shows `LTA-type`=0.397, `LTA-time`=0.280, and `time correlation`=0.477 when $\alpha = 0$, significantly lower than when $\alpha > 0$ (`LTA-type`$>$ 0.5, `LTA-time`$>$ 0.5, and `time correlation`$>$ 0.7). However, the `batch entropy` value remains similar when $\alpha = 0$ (0.422) and $\alpha > 0$ (0.394 on average), meaning modalities are mixed well even if the model does not enforce it. This suggests that `scMultiNODE` can still achieve some degree of integration due to the shared fusion layer between modalities, while the cell type variations and dynamics are not preserved. Nonetheless, the cell integration loss term is essential for learning a joint latent space that effectively captures the diverse variations.

`scMultiNODE` uses the dynamic regularization controlled by $\beta$ to incorporate learned dynamics into the joint latent space. We vary $\beta \in \{0.0, 0.01, 0.1, 1.0, 10.0, 100.0\}$. Supp. Table S9 denotes that removing the dynamic regularization (i.e., $\beta = 0$) results in poor integration where cells cannot be aligned at all (with `batch entropy` close to 0 on both datasets) and cellular dynamics are lost (`time correlation`=0.312 for HC and 0.357 for DR). On adding this regularization (i.e., $\beta > 0$), the joint latent space has much better integration and can learn cellular dynamics to model the development accurately. We also note that a very large $\beta$ may break down the model training and lead to bad performance. For example, `scMultiNODE` 's performance significantly decreases when $\beta = 100$ on the HC dataset. These results imply that the dynamic regularization is essential for aligning modalities and capturing dynamics. Users should select $\beta$ carefully within a reasonable range of $[0.01, 10.0]$.

### S10 Comparison of time costs

We compare the runtime of `scMultiNODE` against the baseline models on the HO dataset, varying the number of cells from 1000 to 6000. We choose HO because it contains the most cells (Supp. Table S1), enabling us to evaluate runtime at scale. All methods are run on an Intel Xeon Platinum 8268 CPU with 32GB of memory. As shown in Supp. Fig. S6, `UnionCom` scales exponentially with the number of cells and `Pamona`'s cost rises sharply on large inputs, whereas `scMultiNODE` scales comparably to most baselines. Thus, despite its additional dynamics-learning step, `scMultiNODE` does not substantially increase computational cost, making it suitable for large-scale temporal multi-modal single-cell datasets.

### S11 Cell Trajectory Analysis

We applied `scMultiNODE` in the cell trajectory analysis on the human cortex (HC) dataset. This dataset contains multifurcating cell trajectories in the human brain cortex, allowing us to validate our analysis with known cell developmental lineages. Apart from baseline multi-modal integration methods, we also compare `scMultiNODE` with static single-modal method AE and a dynamic single-modal method scNODE [35]. scNODE is a novel method that explicitly incorporates cellular dynamics in the latent space. On each dataset, the scNODE's hyperparameters are optimized following the strategy from its original paper.

Moreover, to enable a cross-modal comparison analogous to integration-based methods, we introduce `scNODE (transfer)`, which trains `scNODE` on one modality to infer pseudotime and transfers the predictions to the other modality via regression on the integrated embeddings produced by each baseline integration method. The intuition is that successful integration aligns RNA and ATAC cells in a shared latent space, allowing a regressor trained on source-modality embeddings and pseudotimes to generalize to target-modality embeddings. We evaluate two regressors (k-nearest neighbors and multilayer perceptron) in both transfer directions (RNA-to-ATAC and ATAC-to-RNA) and report the average across directions. The reported `scNODE (transfer)` result is the best performance across all baseline integration methods and regressors, with detailed results provided in Supp. Fig. S15.

Specifically, for each method's latent space, we used Monocle3 [4] to perform trajectory inference and estimate cell pseudotime (Supp. Fig. S8). Monocle3 begins by reducing the dimensionality of the input data (already 50-dimensional latent cell embeddings in our case), followed by the construction of a tree-like principal graph to learn the manifold structure of the data. Branches of the tree correspond to divergent cell fate decisions. To assign pseudotime values, a root node must be selected, which serves as the starting point of the trajectory. We choose the graph node corresponding to the earliest timepoint in the dataset as the root. Monocle3 computes pseudotime as the geodesic distance along the learned principal graph from the root node to each cell, thereby reflecting each cell's relative progression through the underlying biological process. We run Monocle3 following its tutorial (https://cole-trapnell-lab.github.io/monocle3/docs/trajectories/) and use default function parameters. To quantitatively evaluate the accuracy of pseudotime predictions, we calculate the Spearman rank correlation coefficient ($\rho$) between the true cell order labels along the reference trajectory and the inferred pseudotime values. Spearman correlation assesses the degree to which the rank ordering of cells is preserved, making it well-suited for capturing the correctness of developmental progression. A higher correlation value indicates that the inferred pseudotime more accurately reflects the true temporal sequence of cell states.

Additionally, we applied PAGA [31] for pseudotime inference (Supp. Fig. S9), another state-of-the-art trajectory inference method that models cell connectivity by constructing a graph capturing the topology of the data. Pseudotime is then inferred along graph paths, enabling robust trajectory reconstruction across complex lineage structures. Following the recommended workflow (https://scanpy-tutorials.readthedocs.io/en/latest/paga-paul15.html) with default parameters, we again evaluated performance using Spearman correlation (Supp. Fig. S7).

The PAGA-based pseudotime analysis yields conclusions consistent with our main results. On the EN-related trajectory, `scMultiNODE` achieves a Spearman correlation of $\rho = 0.83$, substantially outperforming the strongest integration baseline ($\rho = 0.55$). Similarly, on the IN-mge-related trajectory, `scMultiNODE` reaches $\rho = 0.71$, compared to $\rho = 0.57$ for the best integration baseline. Comparable trends are observed for the IN-cge trajectory. On the oligodendrocyte-related trajectory, `scMultiNODE` attains $\rho = 0.46$, comparable to SCOTv1 ($\rho = 0.52$) while outperforming all other integration baselines.

Compared with single-modality models, `scMultiNODE` consistently outperforms both AE and scNODE on ATAC data across all trajectories. For example, on the EN-related trajectory, AE achieves $\rho = 0.45$, whereas scNODE reaches only $\rho = 0.31$. Overall, these results further demonstrate that `scMultiNODE` faithfully preserves biologically meaningful pseudotime relationships across diverse and complex developmental lineages.

### S12 Identification of Cell Path and Development-Related Genes

We use HC dataset to investigate whether `scMultiNODE` can be used to reconstruct meaningful cell developmental paths and uncover underlying gene programs. The cell path is constructed with the least action path (LAP), which has been used in previous studies [26,25,29] to construct cell fate transitions. The LAP method aims to find the optimal path between two cell states while minimizing their action and transition time. With a little abuse of notation, we let $\mathbf{X}$ denote cell representations in the joint latent space. Specifically, given starting point $\mathbf{X}_0$ and end point $\mathbf{X}_K$, LAP fins a path discretized as a sequence of $K$ points $\mathcal{P} = \{\mathbf{X}_0, \cdots, \mathbf{X}_K\}$. For each segment constrained between $\mathbf{X}_{k-1}$ and $\mathbf{X}_k$, its tangential velocity is defined as $\mathbf{V}_k = \frac{(\mathbf{X}_k - \mathbf{X}_{k-1})}{\varDelta}$ where $\varDelta$ is the timestep taken by cells from $\mathbf{X}_{k-1}$. Therefore, we define the action $\mathcal{S}$ along the path $\mathcal{P}$ as

$$\mathcal{S} = \frac{1}{2}\sum_{k=1}^{K}\Big(\mathbf{V}_k - \text{Drift}_\omega(\widetilde{\mathbf{X}}_k)\Big)^2 \varDelta \quad \text{with } \widetilde{\mathbf{X}}_k = \frac{\mathbf{X}_{k-1} + \mathbf{X} - k}{2}. \tag{S12}$$

Here, LAP method aims to align the tangential velocity $\mathbf{V}_k$ with the differential velocity $\text{Drift}_\omega(\widetilde{\mathbf{X}}_k)$ learned by `scMultiNODE` , while having the least transition time. Therefore, the optimal path is

$$\widehat{\mathcal{P}} \;=\; \underset{\mathcal{P},\varDelta}{\text{argmin}}\, \mathcal{S} \;=\; \underset{\mathcal{P},\varDelta}{\text{argmin}}\, \frac{1}{2}\sum_{k=1}^{K}\Big(\mathbf{V}_k - \text{Drift}(\widetilde{\mathbf{X}}_k, \omega)\Big)^2 \varDelta. \tag{S12}$$

Solving Eq. S12 consists of two iterative steps

(1) Minimize action by fixing path $\mathcal{P}$ and varying the timestep $\varDelta$ through

$$\widehat{\varDelta} = \underset{\varDelta}{\text{argmin}}\, \frac{1}{2}\sum_{k=1}^{K}\left(\frac{\mathbf{X}_k - \mathbf{X}_{k-1}}{\varDelta} - \text{Drift}(\widetilde{\mathbf{X}}_k, \omega)\right)^2 \varDelta. \tag{S12}$$

(2) Minimize action by fixing timestep $\widehat{\varDelta}$ and varying path $\mathcal{P}$

$$\widehat{\mathcal{P}} = \underset{\mathbf{X}_1,\cdots,\mathbf{X}_{K-1}}{\text{argmin}}\, \frac{1}{2}\sum_{k=1}^{K}\left(\frac{\mathbf{X}_k - \mathbf{X}_{k-1}}{\widehat{\varDelta}} - \text{Drift}(\widetilde{\mathbf{X}}_k, \omega)\right)^2 \varDelta. \tag{S12}$$

The starting ($\mathbf{X}_0$) and end point ($\mathbf{X}_K$) are fixed in the optimization.

We use *scipy.optimize.minimize* to solve these two objective functions. In our experiments, we construct two paths from the first timepoint to two cell populations (oligodendrocyte and glutamatergic neuron) with $K = 8$. We set the starting point as the center of cells at the first timepoint ($t = 0$) and the endpoint as the center of the cell population. We initialize timestep as $\varDelta = 1$ and $\mathcal{P}$ as equally spaced points from the starting to end points.

When finding the differentially expressed (DE) genes along the path, we augment the LAP path with its nearest neighbors. Specifically, assuming $\mathcal{P} = \{\mathbf{X}_0, \cdots, \mathbf{X}_K\}$ is the LAP path from $\mathbf{X}_0$ to $\mathbf{X}_K$, we have only eight cells on the path, which is insufficient for DE detection. Therefore, for each $\mathbf{X}_k \in \mathcal{P}$, we find its nearest neighbors in order to augment the path. We use *sklearn.neighbors.NearestNeighbors* to search for ten nearest neighbors. Then, we can use *Scanpy* to detect DE genes for the augmented path with the Wilcoxon rank-sum test. In parallel, we apply *Scanpy* to identify marker genes for each cell type based on either RNA or ATAC data independently (Supp. Table S5). This allows us to compare the DE genes discovered from the `scMultiNODE` joint latent space with traditional marker genes derived from scRNA-seq or scATAC-seq alone. Our results indicate that explicitly modeling cell dynamics in the joint space enables more effective identification of development-associated DE genes compared to single-modal marker gene analysis.

Specifically, we train `scMultiNODE` on the HC dataset, and construct LAP from the starting timepoint ($t = 0$) to oligodendrocytes and glutamatergic neurons (Fig. S11A). Along each trajectory, we apply the Wilcoxon rank-sum test to identify differentially expressed (DE) genes that drive these transitions. Fig. S11B-C plot variance of top-ranked DE genes' expression across timepoints, for cells on the path and out of the path, with randomly selected genes included as controls. Notably, DE genes exhibit stronger trajectory-associated variation than randomly selected controls. These identified DE genes (Supp. Table S5) are well-supported by prior literature. For example, SV2B transcript is expressed in glutamatergic neurons [23]. Also, SOX6 regulates oligodendrocyte proliferation in the central nervous system [15], and SLC1A3 is similarly enriched in oligodendrocytes [19].

Finally, we compare development-related genes identified from our joint latent space with cell type marker genes derived from a standard scRNA-seq/scATAC-seq pipeline (Supp. Table S5). Notably, key developmental genes along the GN and OL trajectories (e.g., SV2B, SOX6) are absent from RNA-derived marker lists despite known functional roles, highlighting limitations of static marker-based analyses. SV2B appears as an ATAC-specific marker, indicating ATAC contributes signals RNA cannot see, which reinforces the value of multi-modal integration for uncovering complementary biological signals.

In summary, `scMultiNODE` provides an interpretable joint latent space that enables not only integration, but also the discovery of development-related, biologically meaningful genes by leveraging complementary information from both modalities.

### S13 Germ Layer Label Transfer

We demonstrate cross-modality germ layer label transfer by leveraging the joint latent space learned by `scMultiNODE` on the unaligned drosophila embryogenesis (DR) dataset. In this setting, germ layer annotations are available only for the RNA modality, and not for the ATAC modality. We assess whether the integrated latent space from each method preserves biologically meaningful structures that enable accurate label transfer from RNA to ATAC.

To evaluate this, we first trained a random forest classifier [18] on the latent representations of RNA cells using their known germ layer labels. The classifier, implemented using the default parameters from the *scikit-learn* package [24], was then applied to the ATAC cell embeddings within the same joint space to infer their germ layer identities. Supp. Fig. S12 indicates that `scMultiNODE` produced coherent ATAC clusters that aligned with known developmental trajectories from broad ectodermal states toward neuroectodermal subpopulations. However, the baseline methods either fail to predict all germ layer labels or do not preserve the underlying developmental trajectories. This implies that the `scMultiNODE` integration captures meaningful biological progression, highlighting its advantage over existing approaches.

Since ground truth germ layer labels for ATAC cells are not available, we validated the predicted labels through two complementary approaches. First, we identified germ layer-specific marker genes from the RNA modality using the Wilcoxon rank-sum test in *Scanpy*, and then examined their expression patterns in the ATAC-derived gene activity matrix, grouped by the predicted germ layer labels. As shown in Supp. Fig. S13, marker gene expression clearly separates the predicted germ layer groups in our model, indicating successful label transfer. In contrast, baseline methods show diffuse or overlapping marker expression across groups, suggesting poor alignment and less effective transfer of germ layer identity. Second, we performed Gene Ontology (GO) enrichment analysis using clusterProfiler [33] on the ATAC-derived gene activity matrix of each predicted germ layer group, focusing on the Biological Process (BP) ontology (Supp. Fig. S14). The enriched GO terms reveal distinct functional profiles for each germ layer group, aligning well with the known biological roles of their respective germ layers.

Table S1: Data descriptions of real-world single-cell datasets used in experiments.

| **ID** | **Dataset** | **Species** | **# cells** (RNA/ATAC) | **# timepoints** (RNA/ATAC) | **Coassay** | **Source** |
|---|---|---|---|---|---|---|
| HC | human cortex | *Homo sapiens* | 2277/2277 | 10/10 | Yes | [36] |
| HO | human organoid | *Homo sapiens* | 10000/10000 | 11/11 | Yes | [13] |
| DR | drosophila embryogenesis | *Drosophila melanogaster* | 2738/4246 | 11/11 | No | [3] |
| MN | mouse neocortex | *Mus musculus* | 6098/1914 | 3/3 | No | [34] |

Table S2: Hyperparameter search space of `scMultiNODE` and baseline methods during hyperparameter tuning.

| **Model** | **Hyperparameters** |
|---|---|
| `scMultiNODE` | number of neighbors $k \in \{5, 10, 25, 50, 75, 100\}$<br>coefficient $\alpha \in \{0.001, 0.01, 0.1, 1.0, 10.0\}$<br>coefficient $\beta \in \{0.001, 0.01, 0.1, 1.0, 10.0\}$ |
| `SCOTv1` | number of neighbors $k \in \{5, 10, 25, 50, 75, 100\}$<br>entropic regularizer coefficient $e \in [0.001, 0.1]$<br>normalize $\in$ {True, False} |
| `SCOTv2` | number of neighbors $k \in \{5, 10, 25, 50, 75, 100\}$<br>entropic regularizer coefficient $eps \in [0.001, 0.1]$<br>marginal relaxation coefficient $\rho \in [0.001, 0.1]$<br>normalize $\in$ {True, False} |
| `UnionCom` | number of neighbors $k \in \{5, 10, 25, 50, 75, 100\}$<br>perplexity $\in \{10, 25, 50, 75, 100\}$<br>$\beta \in \{0.01, 0.1, 1.0, 10.0\}$ |
| `Pamona` | number of neighbors $k \in \{5, 10, 25, 50, 75, 100\}$<br>regularization coefficient epsilon$\in [0.001, 0.1]$<br>trade-off coefficient Lambda $\in \{0.01, 0.1, 1.0, 10.0\}$ |
| `uniPort` | KL coefficient $\in \{0.01, 0.1, 1.0, 10.0\}$<br><br>OT coefficient$\in \{0.01, 0.1, 1.0, 10.0\}$<br>entropy regularization coefficient $\in \{0.01, 0.1, 1.0, 10.0\}$<br>unbalanced OT parameter $\in \{0.01, 0.1, 1.0, 1.0\}$<br>iteration=10000, batch size=32, learning rate=0.0001,<br>diagonal integration mode, MSE loss |
| `Seurat` | number of anchors for CCA $\in \{5, 25, 50, 75, 100, 125, 150, 175, 200\}$<br>number of neighbors when weighting anchors $\in \{5, 25, 50, 75, 100, 125, 150, 175, 200\}$<br>bandwidth of Gaussian kernel $\in \{0.01, 0.05, 0.1, 0.5, 1.0, 5.0, 10.0\}$<br>reference modality $\in$ {RNA, ATAC} |

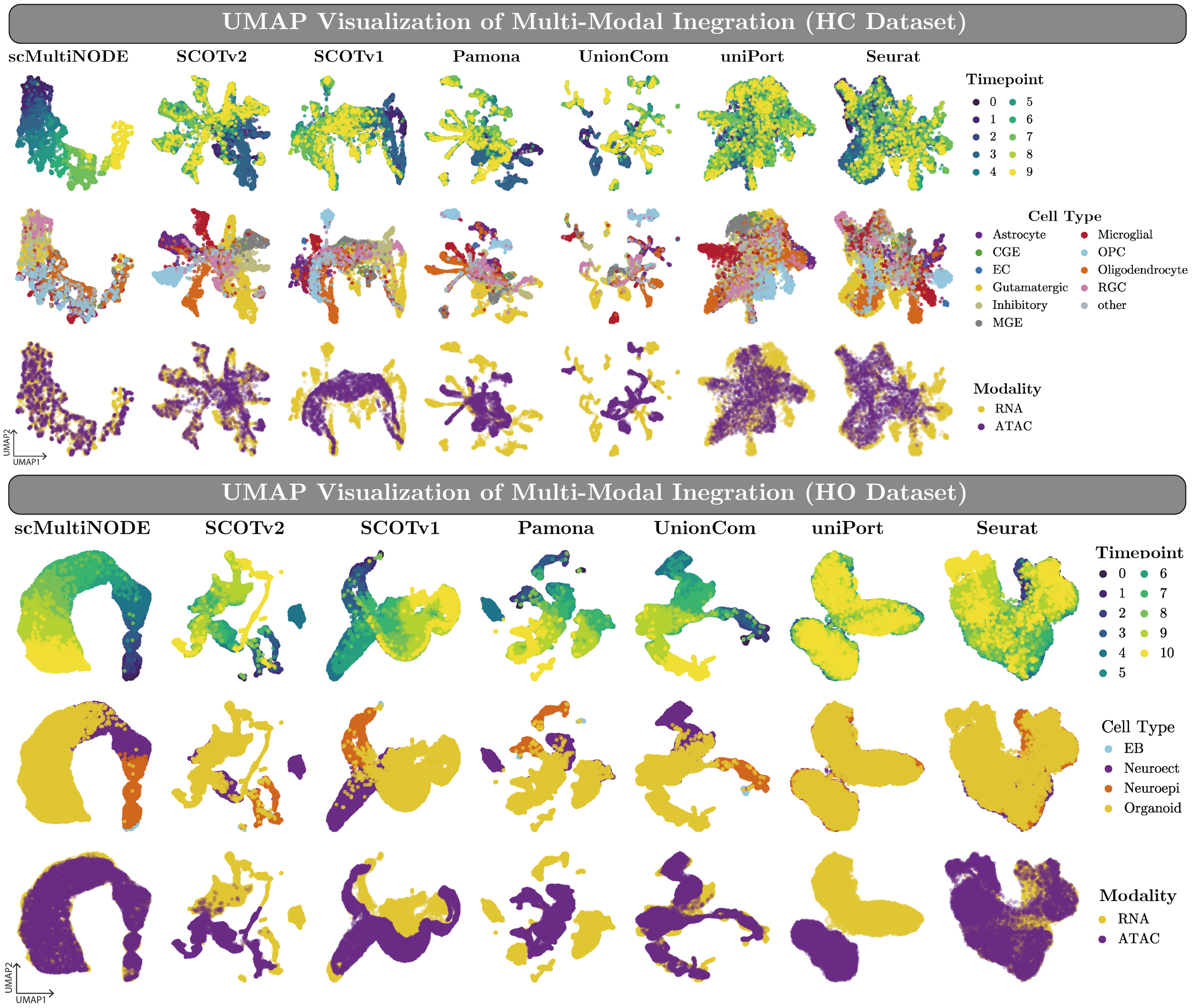


Fig. S1: 2D UMAP visualization of all models' joint latent representations on the coassay datasets. The representations are colored by timepoint labels (*top*), cell types (*middle*), and modality (*bottom*).

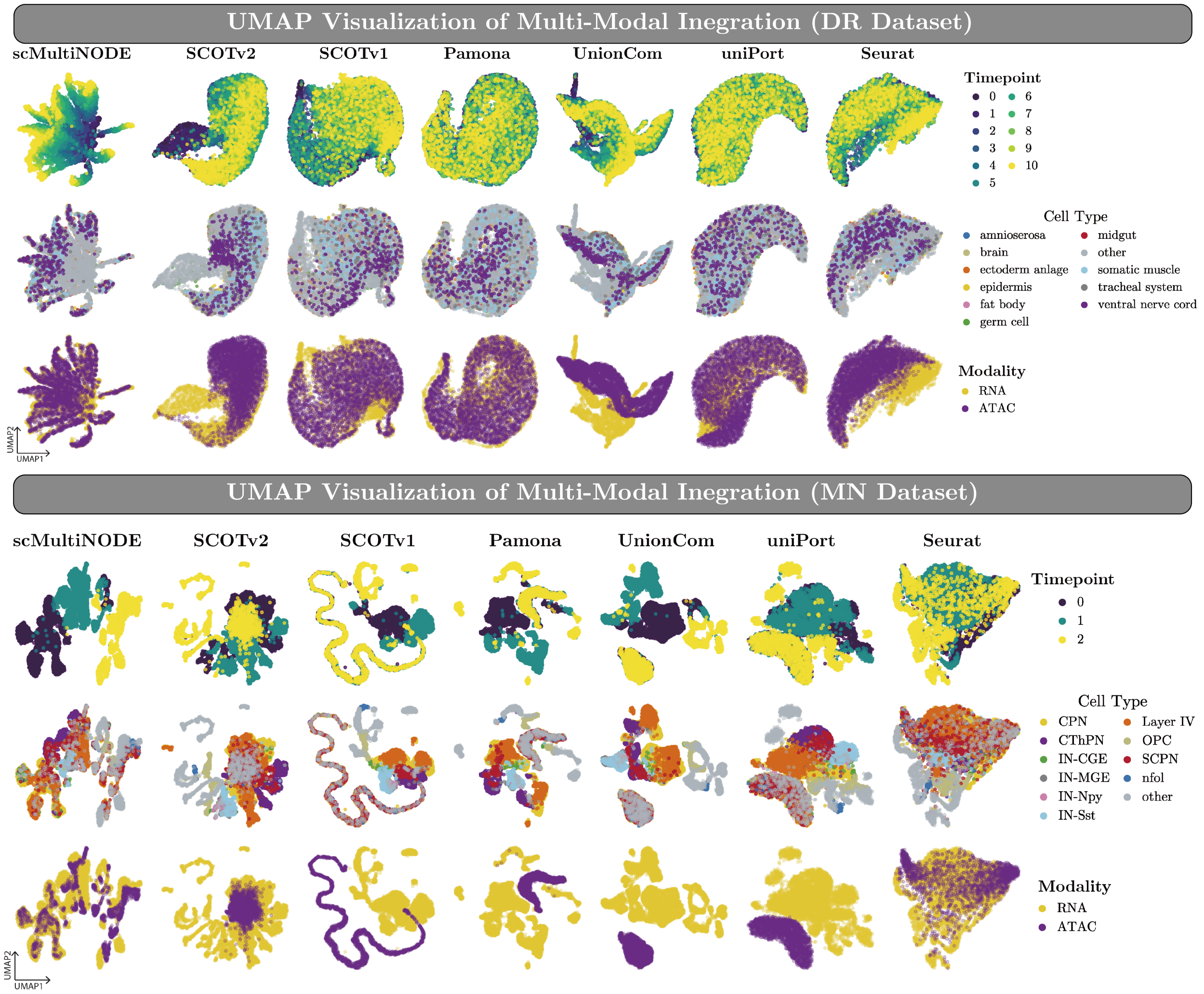


Fig. S2: 2D UMAP visualization of all models' joint latent representations on the unaligned datasets. The representations are colored by timepoint labels (*top*), cell types (*middle*), and modality (*bottom*).

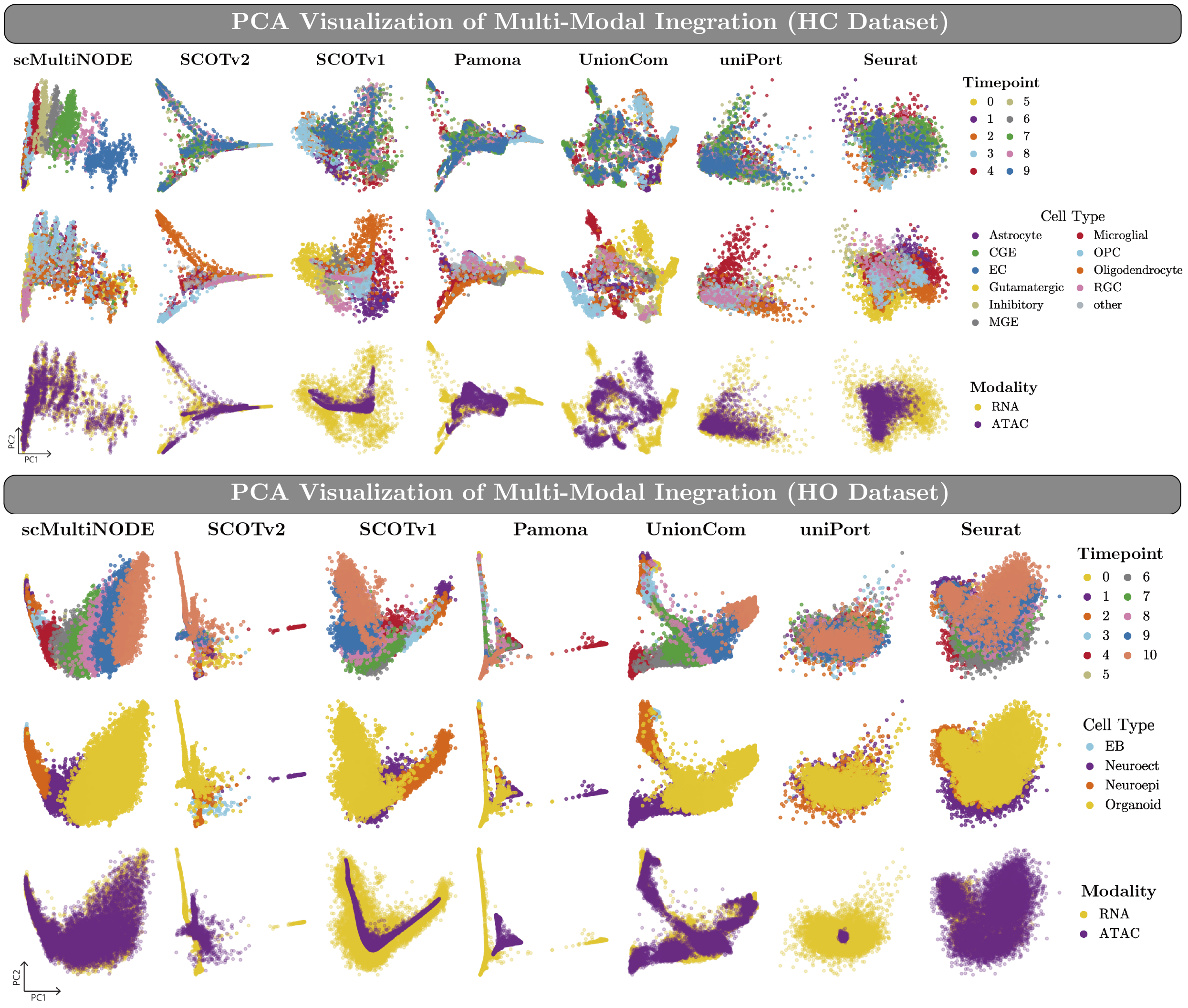


Fig. S3: 2D PCA visualization of all models' joint latent representations on the coassay datasets. The representations are colored by timepoint labels (*top*), cell types (*middle*), and modality (*bottom*).

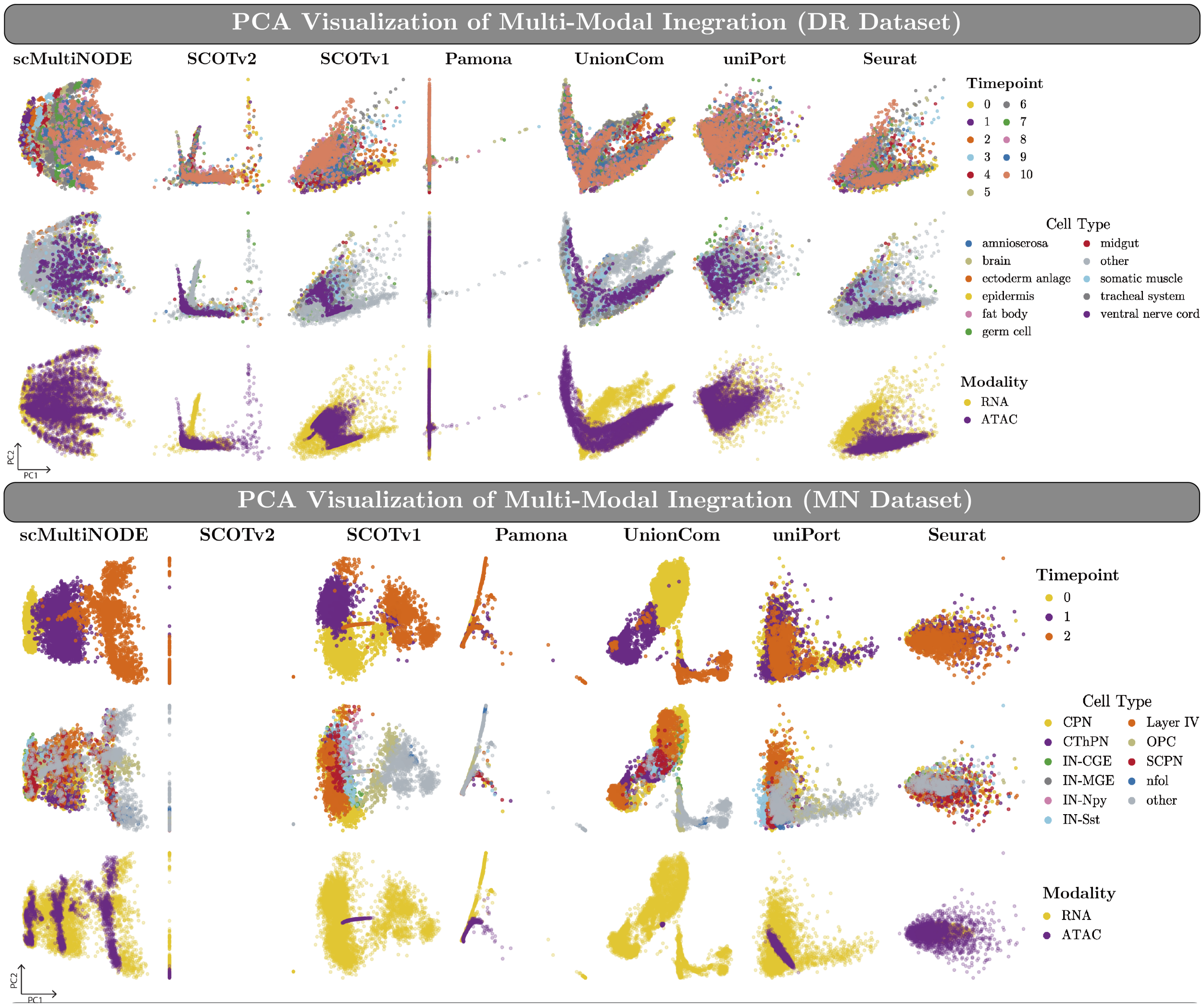


Fig. S4: 2D PCA visualization of all models' joint latent representations on the unaligned datasets. The representations are colored by timepoint labels (*top*), cell types (*middle*), and modality (*bottom*).

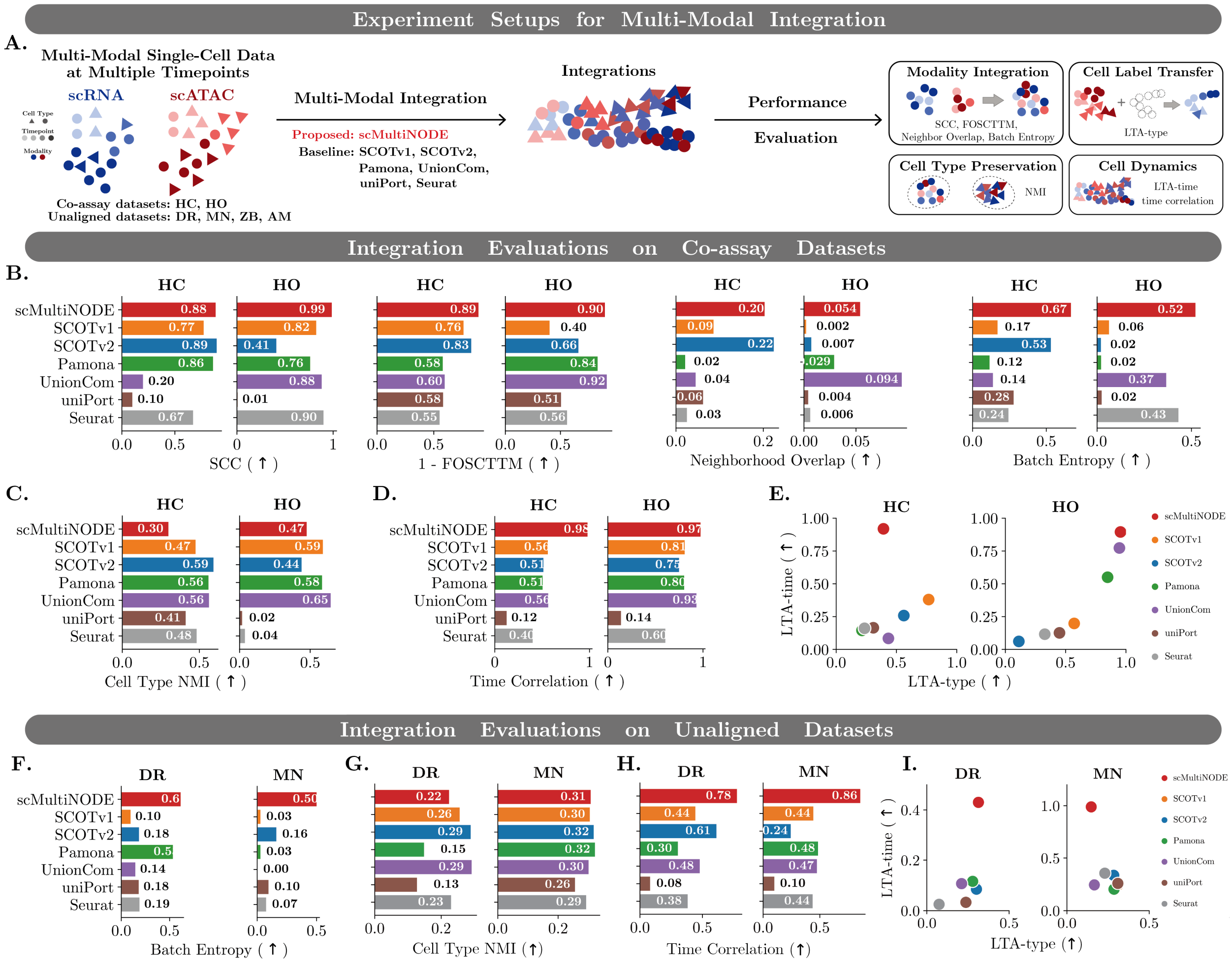


Fig. S5: **Integration evaluation** on two co-assay datasets: HC and HO; two unaligned datasets: DR and MN. (**A**) Schematic overview of integration performance comparison. (**B** and **F**) Performance of modality integration. `Batch entropy` evaluates the integration of all datasets. Three metrics that are only available on co-assay datasets: Spearman correlation coefficient (`SCC`), `neighborhood overlap`, and fraction of samples closer than the true match (`FOSCTTM`). We show `1-FOSCTTM` instead of `FOSCTTM` here to unify figure plotting for all metrics, where a higher metric value implies better integration performance. (**C** and **G**) Cell type conservation after integration. We calculate the Normalized Mutual Information (`NMI`) score, with higher values indicating better preservation of cell groups. (**D** and **H**) `Time correlation` evaluates integrations on capturing cell dynamics. (**E** and **I**) Label transfer accuracy (LTA) across modalities. We plot the `LTA-type` and `LTA-time` scores on the X and Y axes for all the methods, respectively. `LTA-type` measures cell type variations and `LTA-time` for temporal variations. `scMultiNODE` outperforms existing approaches in integrating temporal multi-modal single-cell data, demonstrating strong performance on both co-assay and unaligned datasets while preserving cell type variation and underlying cellular dynamics.

Table S3: Evaluation of model integration on two co-assay datasets, HC and HO. The **red bold** and blue underlined numbers indicate the best and the second best performance, respectively.

| **Method** (HC) | **Modality Integration** | | | | **Cell Label Transfer** | **Cell Type Preservation** | **Cellular Dynamic** | |
|---|---|---|---|---|---|---|---|---|
| | Batch Entropy (↑) | FOSCTTM (↓) | Neighborhood Overlap (↑) | SCC (↑) | LTA-type (↑) | NMI (↑) | LTA-time (↑) | Time Correlation (↑) |
| `scMultiNODE` | **0.667** | **0.106** | 0.203 | 0.884 | 0.392 | 0.299 | **0.919** | **0.979** |
| `SCOTv1` | 0.169 | 0.238 | 0.086 | 0.771 | 0.561 | 0.475 | 0.258 | 0.560 |
| `SCOTv2` | 0.531 | 0.170 | **0.224** | **0.894** | **0.767** | **0.591** | 0.380 | 0.513 |
| `Pamona` | 0.115 | 0.421 | 0.021 | 0.859 | 0.214 | 0.559 | 0.145 | 0.505 |
| `UnionCom` | 0.138 | 0.404 | 0.045 | 0.199 | 0.433 | 0.562 | 0.084 | 0.564 |
| `uniPort` | 0.278 | 0.418 | 0.062 | 0.098 | 0.308 | 0.411 | 0.165 | 0.125 |
| `Seurat` | 0.243 | 0.449 | 0.025 | 0.671 | 0.235 | 0.482 | 0.161 | 0.400 |

| **Method** (HO) | **Modality Integration** | | | | **Cell Label Transfer** | **Cell Type Preservation** | **Cellular Dynamic** | |
|---|---|---|---|---|---|---|---|---|
| | Batch Entropy (↑) | FOSCTTM (↓) | Neighborhood Overlap (↑) | SCC (↑) | LTA-type (↑) | NMI (↑) | LTA-time (↑) | Time Correlation (↑) |
| `scMultiNODE` | **0.521** | 0.097 | **0.0544** | **0.986** | **0.955** | 0.475 | **0.895** | **0.974** |
| `SCOTv1` | 0.063 | 0.599 | 0.0019 | 0.824 | 0.112 | 0.590 | 0.061 | 0.807 |
| `SCOTv2` | 0.020 | 0.337 | 0.0069 | 0.409 | 0.571 | 0.439 | 0.198 | 0.748 |
| `Pamona` | 0.021 | 0.163 | 0.0291 | 0.761 | 0.848 | 0.584 | 0.550 | 0.802 |
| `UnionCom` | 0.366 | **0.080** | 0.094 | 0.881 | 0.947 | **0.645** | 0.773 | 0.931 |
| `uniPort` | 0.024 | 0.495 | 0.0042 | 0.007 | 0.449 | 0.018 | 0.126 | 0.138 |
| `Seurat` | 0.431 | 0.440 | 0.0061 | 0.900 | 0.327 | 0.038 | 0.116 | 0.604 |

Table S4: Evaluation of model integration on two unaligned datasets DR and MN. The **red bold** and blue underlined numbers indicate the best and the second best performance, respectively. Because there is no cell-to-cell correspondence for unaligned datasets, we remove `FOSCTTM`, `neighborhood overlap`, and `SCC` here, which require such information.

| **Method** (DR) | **Modality Integration** | **Cell Label Transfer** | **Cell Type Preservation** | **Cellular Dynamic** | |
|---|---|---|---|---|---|
| | Batch Entropy (↑) | LTA-type (↑) | NMI (↑) | LTA-time (↑) | Time Correlation (↑) |
| `scMultiNODE` | **0.614** | **0.314** | 0.224 | **0.430** | **0.777** |
| `SCOTv1` | 0.096 | 0.297 | 0.256 | 0.094 | 0.443 |
| `SCOTv2` | 0.183 | 0.302 | 0.289 | 0.085 | 0.613 |
| `Pamona` | 0.534 | 0.279 | 0.148 | 0.116 | 0.303 |
| `UnionCom` | 0.145 | 0.212 | **0.292** | 0.107 | 0.478 |
| `uniPort` | 0.180 | 0.238 | 0.127 | 0.033 | 0.081 |
| `Seurat` | 0.188 | 0.074 | 0.230 | 0.025 | 0.381 |

| **Method** (MN) | **Modality Integration** | **Cell Label Transfer** | **Cell Type Preservation** | **Cellular Dynamic** | |
|---|---|---|---|---|---|
| | Batch Entropy (↑) | LTA-type (↑) | NMI (↑) | LTA-time (↑) | Time Correlation (↑) |
| `scMultiNODE` | **0.500** | 0.148 | 0.308 | **0.989** | **0.856** |
| `SCOTv1` | 0.027 | **0.311** | 0.305 | 0.245 | 0.442 |
| `SCOTv2` | 0.161 | 0.285 | 0.318 | 0.336 | 0.243 |
| `Pamona` | 0.027 | 0.287 | **0.322** | 0.204 | 0.484 |
| `UnionCom` | 0.001 | 0.167 | 0.301 | 0.245 | 0.474 |
| `uniPort` | 0.095 | 0.310 | 0.256 | 0.259 | 0.099 |
| `Seurat` | 0.075 | 0.231 | 0.293 | 0.356 | 0.436 |

Table S5: The top 10 DE genes of oligodendrocyte (OL) and glutamatergic neuron (GN) paths, obtained from `scMultiNODE` 's joint latent space. We also show the top 10 marker genes of the OL and GN cell types, derived from RNA and ATAC data.

| | | |
|---|---|---|
| DE genes found in `scMultiNODE` joint latent space | OL path | SOX6, SLC1A3, NEAT1, ADGRV1, SLC1A2, PRKG1, GPC5, GLUL, SFMBT2, ATP1A2 |
| | GN path | SV2B, ANO3, MTUS2, ZNF536, PDE8B, BMPER, ENSG00000251680, KIRREL3, FSTL5, SEC14L5, GRIN2A, CLMN |
| RNA-derived marker genes | OL cell type | CTNNA3, SLC24A2, ST18, RNF220, PLP1, PIP4K2A, MAP7, MBP, DOCK10, MOBP |
| | GN cell type | SATB2, RBFOX1, NRG1, ROBO2, RALYL, MIR137HG, KCNQ5, IQCJ-SCHIP1, DLGAP2, RYR2 |
| ATAC-derived marker genes | OL cell type | RNF220, POLR2F, TFEB, FA2H, AATK, FAM102A, C10orf90, PRIMA1, CLMN, BCAR1 |
| | GN cell type | RBFOX1, SATB2, NELL2, EFCAB6, MYT1L, MPPED1, NKAIN2, SV2B, ROBO2, SLC44A5 |

Table S6: `scMultiNODE` integration performance when using different joint latent space size $d$.

**HC Dataset**

| **Latent Size ($d$)** | **Modality Integration** | | | | **Cell-Type Variation** | **Cellular Dynamic** | |
|---|---|---|---|---|---|---|---|
| | Batch Entropy (↑) | FOSCTTM (↓) | Neighborhood Overlap (↑) | SCC (↑) | LTA-type (↑) | LTA-time (↑) | Time Correlation (↑) |
| 10 | 0.575 | 0.324 | 0.083 | 0.921 | 0.316 | 0.434 | 0.674 |
| 50 | 0.643 | 0.108 | 0.183 | 0.923 | 0.415 | 0.803 | 0.944 |
| 100 | 0.647 | 0.094 | 0.232 | 0.853 | 0.441 | 0.864 | 0.965 |
| 150 | 0.683 | 0.127 | 0.178 | 0.863 | 0.375 | 0.885 | 0.959 |
| 200 | 0.677 | 0.131 | 0.152 | 0.841 | 0.360 | 0.886 | 0.972 |

**DR Dataset**

| **Latent Size ($d$)** | **Modality Integration** | **Cell-Type Variation** | **Cellular Dynamic** | |
|---|---|---|---|---|
| | Batch Entropy (↑) | LTA-type (↑) | LTA-time (↑) | Time Correlation (↑) |
| 10 | 0.589 | 0.333 | 0.462 | 0.734 |
| 50 | 0.520 | 0.315 | 0.395 | 0.635 |
| 100 | 0.483 | 0.341 | 0.403 | 0.552 |
| 150 | 0.459 | 0.390 | 0.408 | 0.614 |
| 200 | 0.545 | 0.308 | 0.441 | 0.695 |

Table S7: `scMultiNODE` integration performance when using different number of neighbors ($k$) in GW optimal transport.

**HC Dataset**

| **Number of Neighbors ($k$)** | **Modality Integration** | | | | **Cell-Type Variation** | **Cellular Dynamic** | |
|---|---|---|---|---|---|---|---|
| | Batch Entropy (↑) | FOSCTTM (↓) | Neighborhood Overlap (↑) | SCC (↑) | LTA-type (↑) | LTA-time (↑) | Time Correlation (↑) |
| 5 | 0.657 | 0.124 | 0.162 | 0.872 | 0.354 | 0.826 | 0.971 |
| 10 | 0.650 | 0.180 | 0.157 | 0.881 | 0.493 | 0.691 | 0.881 |
| 50 | 0.647 | 0.152 | 0.159 | 0.911 | 0.432 | 0.739 | 0.939 |
| 100 | 0.651 | 0.130 | 0.174 | 0.870 | 0.399 | 0.837 | 0.933 |
| 150 | 0.620 | 0.153 | 0.179 | 0.906 | 0.360 | 0.759 | 0.948 |
| 200 | 0.646 | 0.136 | 0.133 | 0.921 | 0.326 | 0.778 | 0.957 |
| Best Baseline | 0.531 | 0.170 | 0.224 | 0.894 | 0.767 | 0.380 | 0.564 |

**DR Dataset**

| **Number of Neighbors ($k$)** | **Modality Integration** | **Cell-Type Variation** | **Cellular Dynamic** | |
|---|---|---|---|---|
| | Batch Entropy (↑) | LTA-type (↑) | LTA-time (↑) | Time Correlation (↑) |
| 5 | 0.591 | 0.436 | 0.571 | 0.779 |
| 10 | 0.613 | 0.372 | 0.430 | 0.777 |
| 50 | 0.598 | 0.335 | 0.587 | 0.771 |
| 100 | 0.615 | 0.350 | 0.537 | 0.760 |
| 150 | 0.534 | 0.399 | 0.544 | 0.768 |
| 200 | 0.611 | 0.324 | 0.571 | 0.766 |
| Best Baseline | 0.534 | 0.116 | 0.302 | 0.613 |

Table S8: `scMultiNODE` integration performance when using different fusion coefficient ($\alpha$) in Eq. 5.

**HC Dataset**

| **Fusion Coefficient** ($\alpha$) | **Modality Integration** | | | | **Cell-Type Variation** | **Cellular Dynamic** | |
|---|---|---|---|---|---|---|---|
| | Batch Entropy (↑) | FOSCTTM (↓) | Neighborhood Overlap (↑) | SCC (↑) | LTA-type (↑) | LTA-time (↑) | Time Correlation (↑) |
| 0.0 | 0.624 | 0.259 | 0.113 | 0.883 | 0.369 | 0.451 | 0.811 |
| 0.01 | 0.659 | 0.222 | 0.132 | 0.841 | 0.379 | 0.629 | 0.891 |
| 0.1 | 0.657 | 0.115 | 0.170 | 0.911 | 0.400 | 0.788 | 0.965 |
| 1.0 | 0.659 | 0.144 | 0.153 | 0.903 | 0.277 | 0.879 | 0.965 |
| 10.0 | 0.658 | 0.097 | 0.196 | 0.939 | 0.394 | 0.894 | 0.976 |
| 100.0 | 0.640 | 0.329 | 0.075 | 0.754 | 0.216 | 0.662 | 0.816 |
| Best Baseline | 0.531 | 0.170 | 0.224 | 0.894 | 0.767 | 0.380 | 0.564 |

**DR Dataset**

| **Fusion Coefficient** ($\alpha$) | **Modality Integration** | **Cell-Type Variation** | **Cellular Dynamic** | |
|---|---|---|---|---|
| | Batch Entropy (↑) | LTA-type (↑) | LTA-time (↑) | Time Correlation (↑) |
| 0.0 | 0.422 | 0.397 | 0.280 | 0.477 |
| 0.01 | 0.385 | 0.548 | 0.548 | 0.724 |
| 0.1 | 0.444 | 0.575 | 0.575 | 0.774 |
| 1.0 | 0.394 | 0.517 | 0.517 | 0.753 |
| 10.0 | 0.385 | 0.505 | 0.505 | 0.719 |
| 100.0 | 0.363 | 0.601 | 0.601 | 0.782 |
| Best Baseline | 0.534 | 0.116 | 0.302 | 0.613 |

Table S9: `scMultiNODE` integration performance when using different dynamic regularization coefficient ($\beta$) in Eq. 10.

**HC Dataset**

| **Dynamic Regularization Coefficient** ($\beta$) | **Modality Integration** | | | | **Cell-Type Variation** | **Cellular Dynamic** | |
|---|---|---|---|---|---|---|---|
| | Batch Entropy (↑) | FOSCTTM (↓) | Neighborhood Overlap (↑) | SCC (↑) | LTA-type (↑) | LTA-time (↑) | Time Correlation (↑) |
| 0.0 | 0.003 | 0.494 | 0.018 | 0.312 | 0.105 | 0.110 | 0.312 |
| 0.01 | 0.621 | 0.151 | 0.129 | 0.947 | 0.326 | 0.767 | 0.947 |
| 0.1 | 0.648 | 0.118 | 0.164 | 0.942 | 0.393 | 0.807 | 0.942 |
| 1.0 | 0.669 | 0.160 | 0.168 | 0.939 | 0.355 | 0.831 | 0.939 |
| 10.0 | 0.680 | 0.207 | 0.123 | 0.931 | 0.329 | 0.863 | 0.931 |
| 100.0 | 0.669 | 0.286 | 0.101 | 0.775 | 0.383 | 0.604 | 0.775 |
| Best Baseline | 0.531 | 0.170 | 0.224 | 0.894 | 0.767 | 0.380 | 0.564 |

**DR Dataset**

| **Dynamic Regularization Coefficient** ($\beta$) | **Modality Integration** | **Cell-Type Variation** | **Cellular Dynamic** | |
|---|---|---|---|---|
| | Batch Entropy (↑) | LTA-type (↑) | LTA-time (↑) | Time Correlation (↑) |
| 0.0 | 0.002 | 0.335 | 0.124 | 0.357 |
| 0.01 | 0.454 | 0.271 | 0.339 | 0.752 |
| 0.1 | 0.392 | 0.427 | 0.479 | 0.761 |
| 1.0 | 0.558 | 0.284 | 0.447 | 0.611 |
| 10.0 | 0.262 | 0.319 | 0.246 | 0.414 |
| 100.0 | 0.597 | 0.396 | 0.459 | 0.567 |
| Best Baseline | 0.534 | 0.116 | 0.302 | 0.613 |

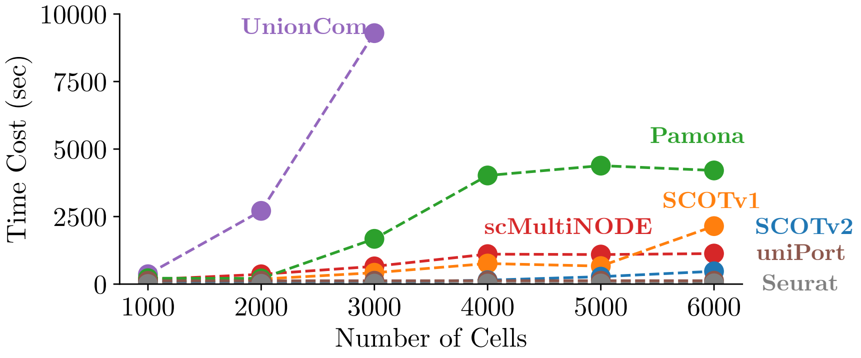


Fig. S6: Runtime comparison of `scMultiNODE` and baselines as the number of cells increases, evaluated on the HO dataset.

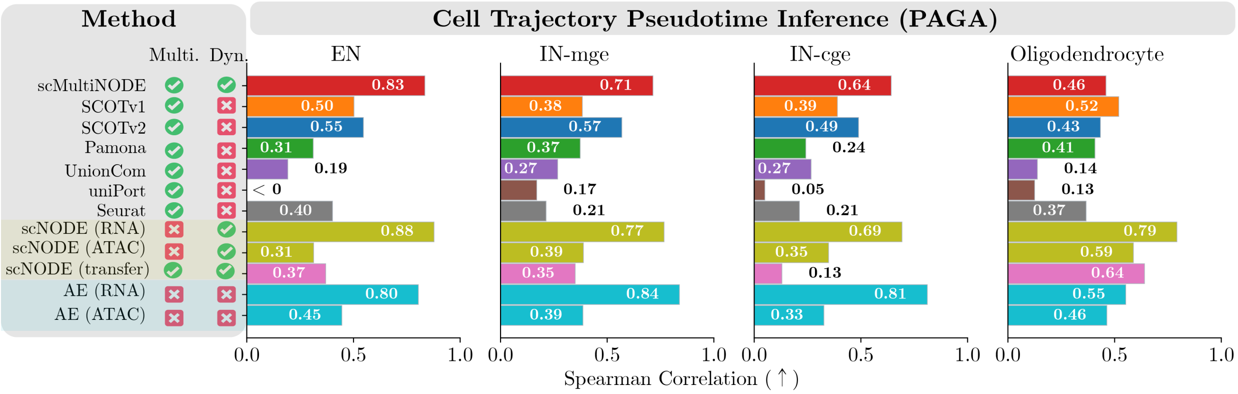


Fig. S7: Spearman correlation coefficient ($\rho$) between ground truth cell state orders and PAGA inferred pseudotime of the integration ("Multi." indicates multi-modal integration method, "Dyn." denotes models incorporating cell dynamics).

Table S10: Evaluation of `scMultiNODE` 's integration on two co-assay datasets, learned with and without cell type information. The **red bold** numbers indicate the better `scMultiNODE` variation.

| **Method** (HC) | **Modality Integration** | | | | **Cell Label Transfer** | **Cell Type Preservation** | **Cellular Dynamic** | |
|---|---|---|---|---|---|---|---|---|
| | Batch Entropy (↑) | FOSCTTM (↓) | Neighborhood Overlap (↑) | SCC (↑) | LTA-type (↑) | NMI (↑) | LTA-time (↑) | Time Correlation (↑) |
| `scMultiNODE` w/o supervision | **0.667** | **0.106** | 0.203 | **0.884** | 0.392 | 0.299 | **0.919** | **0.979** |
| `scMultiNODE` w/ supervision | 0.453 | 0.143 | **0.258** | 0.867 | **0.690** | **0.614** | 0.515 | 0.911 |
| Best baseline | 0.531 | 0.170 | 0.224 | 0.894 | 0.767 | 0.591 | 0.380 | 0.564 |

| **Method** (HO) | **Modality Integration** | | | | **Cell Label Transfer** | **Cell Type Preservation** | **Cellular Dynamic** | |
|---|---|---|---|---|---|---|---|---|
| | Batch Entropy (↑) | FOSCTTM (↓) | Neighborhood Overlap (↑) | SCC (↑) | LTA-type (↑) | NMI (↑) | LTA-time (↑) | Time Correlation (↑) |
| `scMultiNODE` w/o supervision | **0.521** | **0.097** | **0.054** | **0.986** | 0.955 | 0.475 | **0.895** | **0.974** |
| `scMultiNODE` w/ supervision | 0.221 | 0.134 | 0.040 | 0.975 | **0.969** | **0.538** | 0.731 | 0.948 |
| Best Baseline | 0.431 | 0.080 | 0.029 | 0.900 | 0.947 | 0.645 | 0.773 | 0.931 |

Table S11: Evaluation of `scMultiNODE` 's integration on unaligned datasets, learned with and without cell type information. The **red bold** numbers indicate the better `scMultiNODE` variation.

| **Method** (DR) | **Modality Integration** | **Cell Label Transfer** | **Cell Type Preservation** | **Cellular Dynamic** | |
|---|---|---|---|---|---|
| | Batch Entropy (↑) | LTA-type (↑) | NMI (↑) | LTA-time (↑) | Time Correlation (↑) |
| `scMultiNODE` w/o supervision | **0.614** | 0.314 | 0.224 | **0.430** | 0.777 |
| `scMultiNODE` w/ supervision | 0.175 | **0.789** | **0.625** | 0.341 | **0.859** |
| Best Baseline | 0.534 | 0.302 | 0.292 | 0.116 | 0.613 |

| **Method** (MN) | **Modality Integration** | **Cell Label Transfer** | **Cell Type Preservation** | **Cellular Dynamic** | |
|---|---|---|---|---|---|
| | Batch Entropy (↑) | LTA-type (↑) | NMI (↑) | LTA-time (↑) | Time Correlation (↑) |
| `scMultiNODE` w/o supervision | **0.500** | 0.148 | 0.308 | **0.989** | **0.856** |
| `scMultiNODE` w/ supervision | 0.078 | **0.476** | **0.696** | 0.699 | 0.736 |
| Best Baseline | 0.161 | 0.311 | 0.322 | 0.356 | 0.484 |

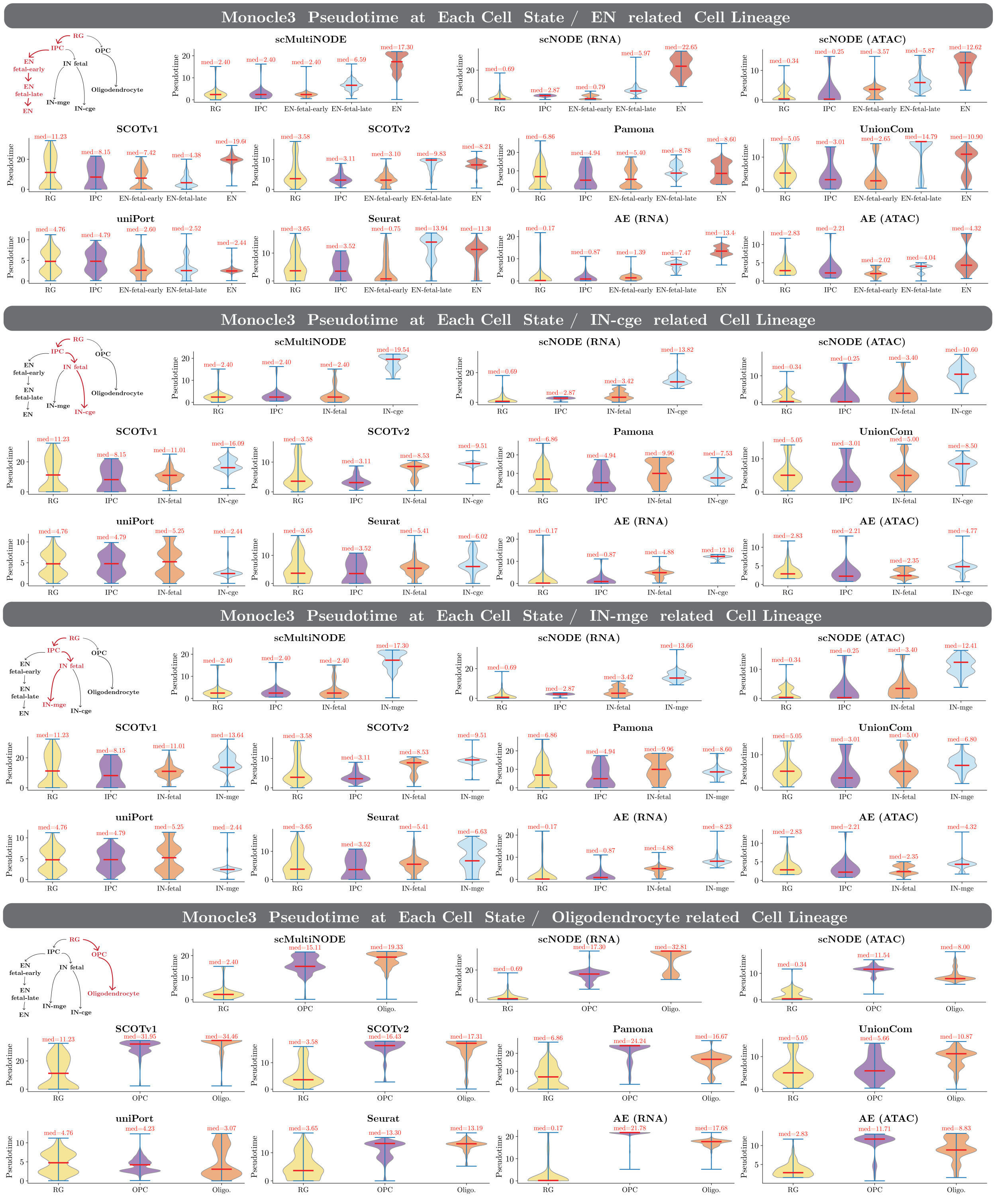


Fig. S8: Violin plots showing Monocle3 pseudotime distributions inferred from the integrated latent space of `scMultiNODE` and baseline integration methods, as well as single-modality models (AE and scNODE). Cells are grouped by annotated cell states, and median pseudotime values ("med") are indicated for each group. The trajectory originates from RG and branches into four lineages (RG: Radial Glia, IPC: Intermediate Progenitor Cells, OPC: Oligodendrocyte Precursor Cells, EN: Excitatory Neurons, IN: Inhibitory Neurons).

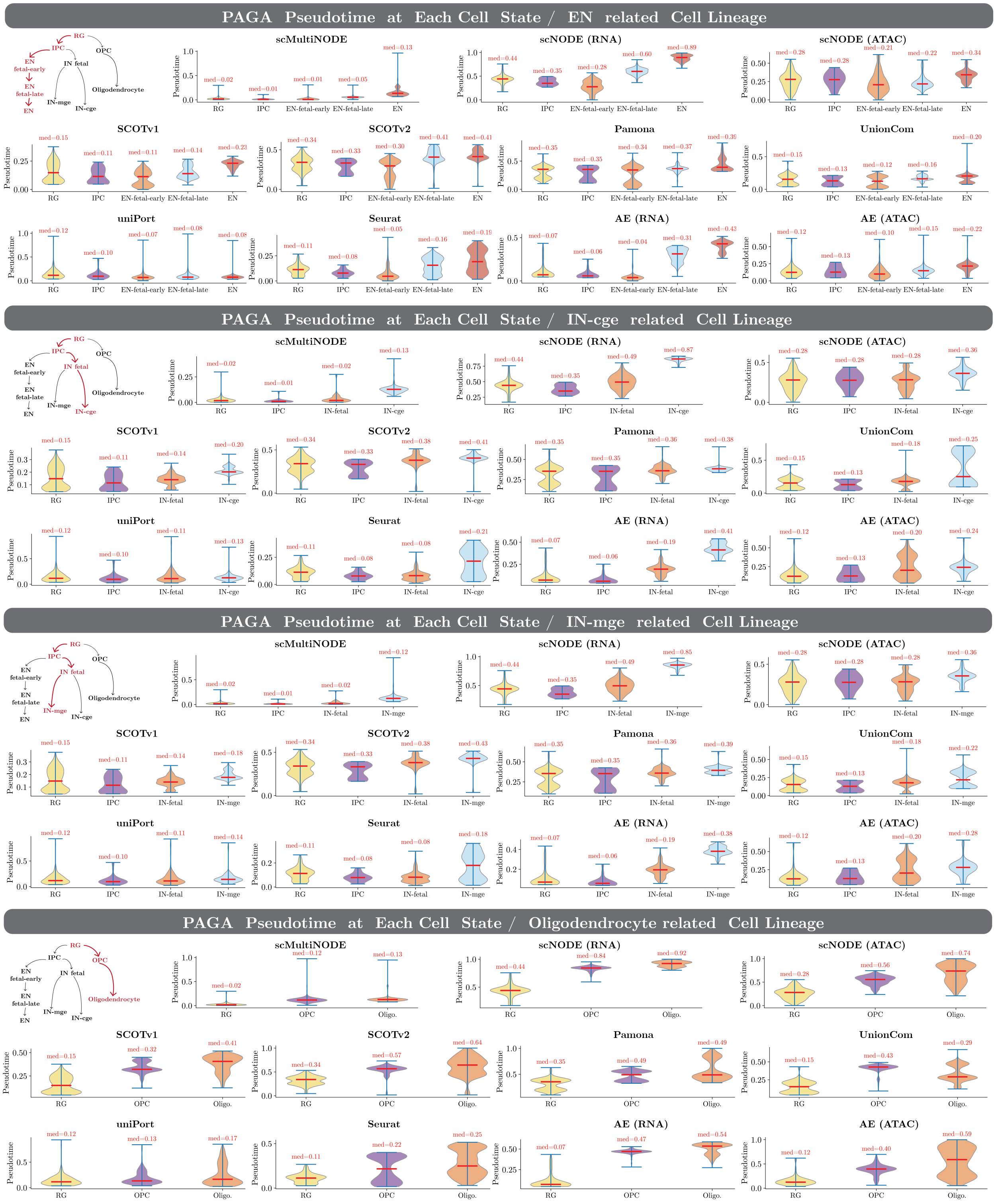


Fig. S9: Violin plots showing PAGA pseudotime distributions inferred from the integrated latent space of `scMultiNODE` and baseline integration methods, as well as single-modality models (AE and scNODE). Cells are grouped by annotated cell states, and median pseudotime values ("med") are indicated for each group. The trajectory originates from RG and branches into four lineages (RG: Radial Glia, IPC: Intermediate Progenitor Cells, OPC: Oligodendrocyte Precursor Cells, EN: Excitatory Neurons, IN: Inhibitory Neurons).

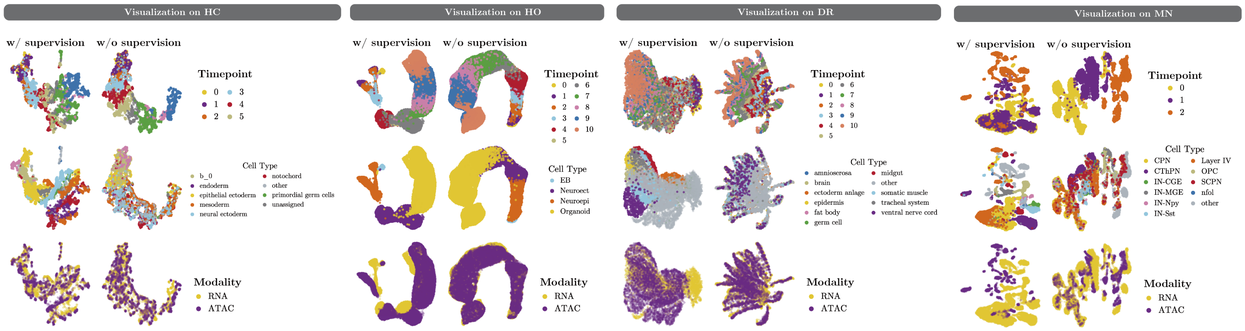

Fig. S10: 2D UMAP visualization of `scMultiNODE` 's joint latent representations, learned with and without cell type supervisions. The representations are colored by timepoint labels (*top*), cell types (*middle*), and modality (*bottom*).

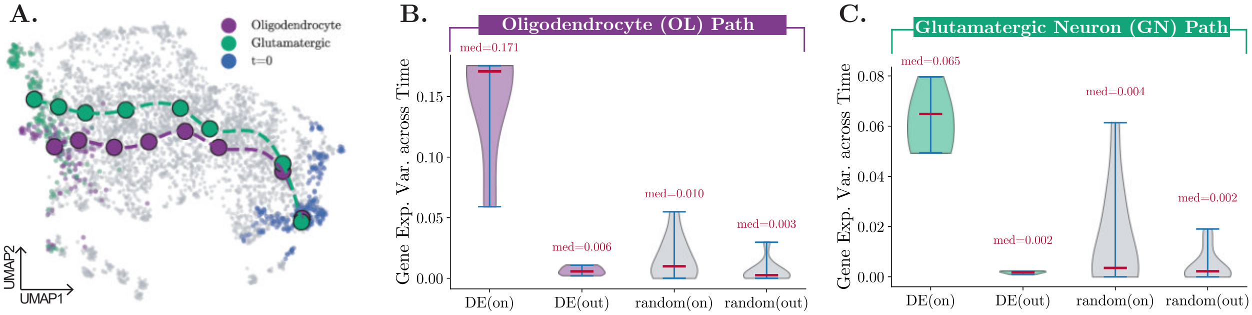


Fig. S11: (**A**) 2D UMAP visualization of the least action path from cells at the starting point ($t = 0$) to the oligodendrocyte (OL) or glutamatergic neuron (GN) populations. (**C**, **C**) Variance of gene expression across timepoints for top-ranked DE genes. We plot the gene expression temporal variance of the top five DE genes for cells on the path (`DE(on)`) and out of the path (`DE(out)`). We also show the temporal variance of five random genes for cells on the path (`random(on)`) and out of the path (`random(out)`).

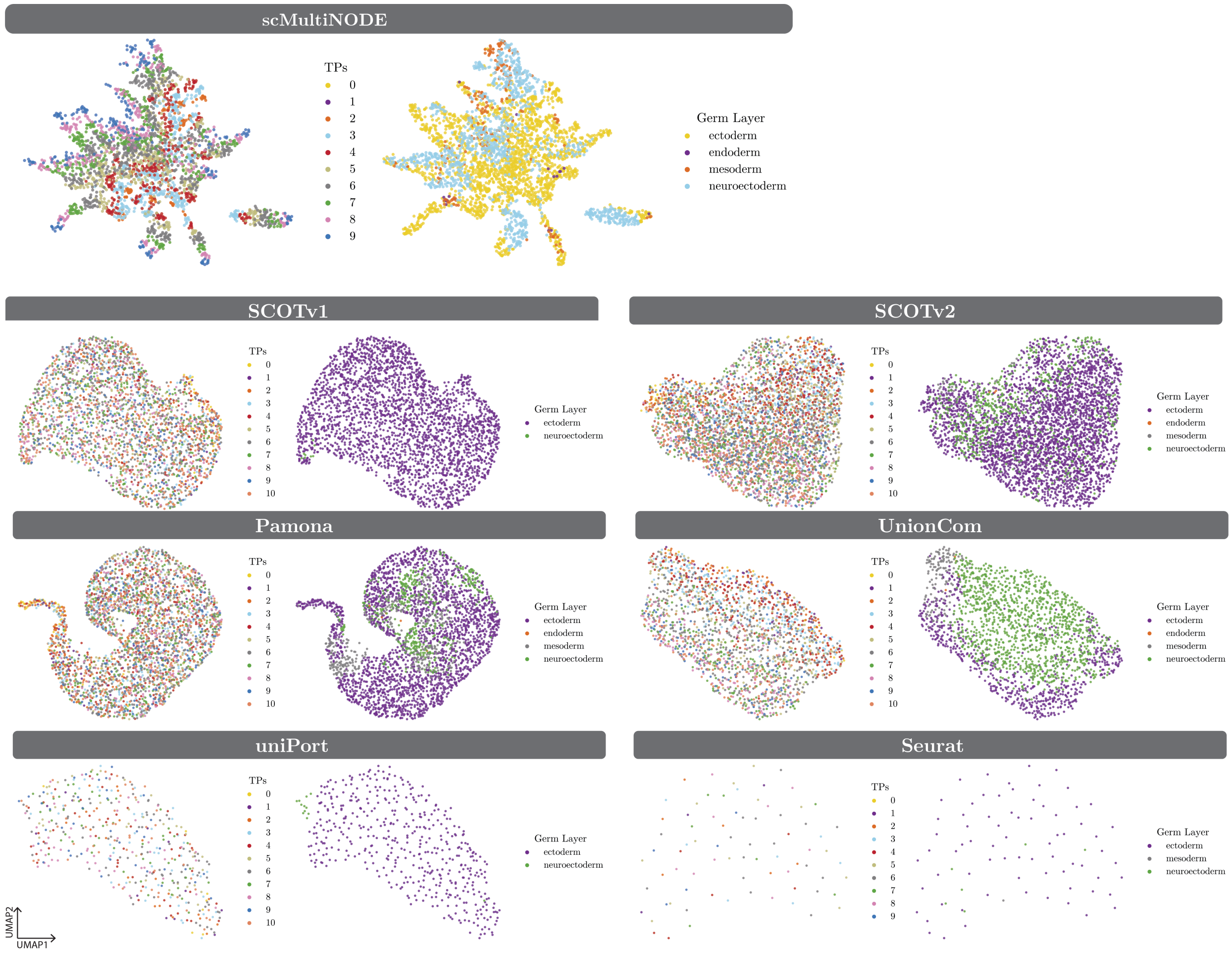


Fig. S12: 2D UMAP visualization of ATAC cells from scMultiNODE and baseline methods' integrations, with cells colored by timepoints and predicted germ layer labels.

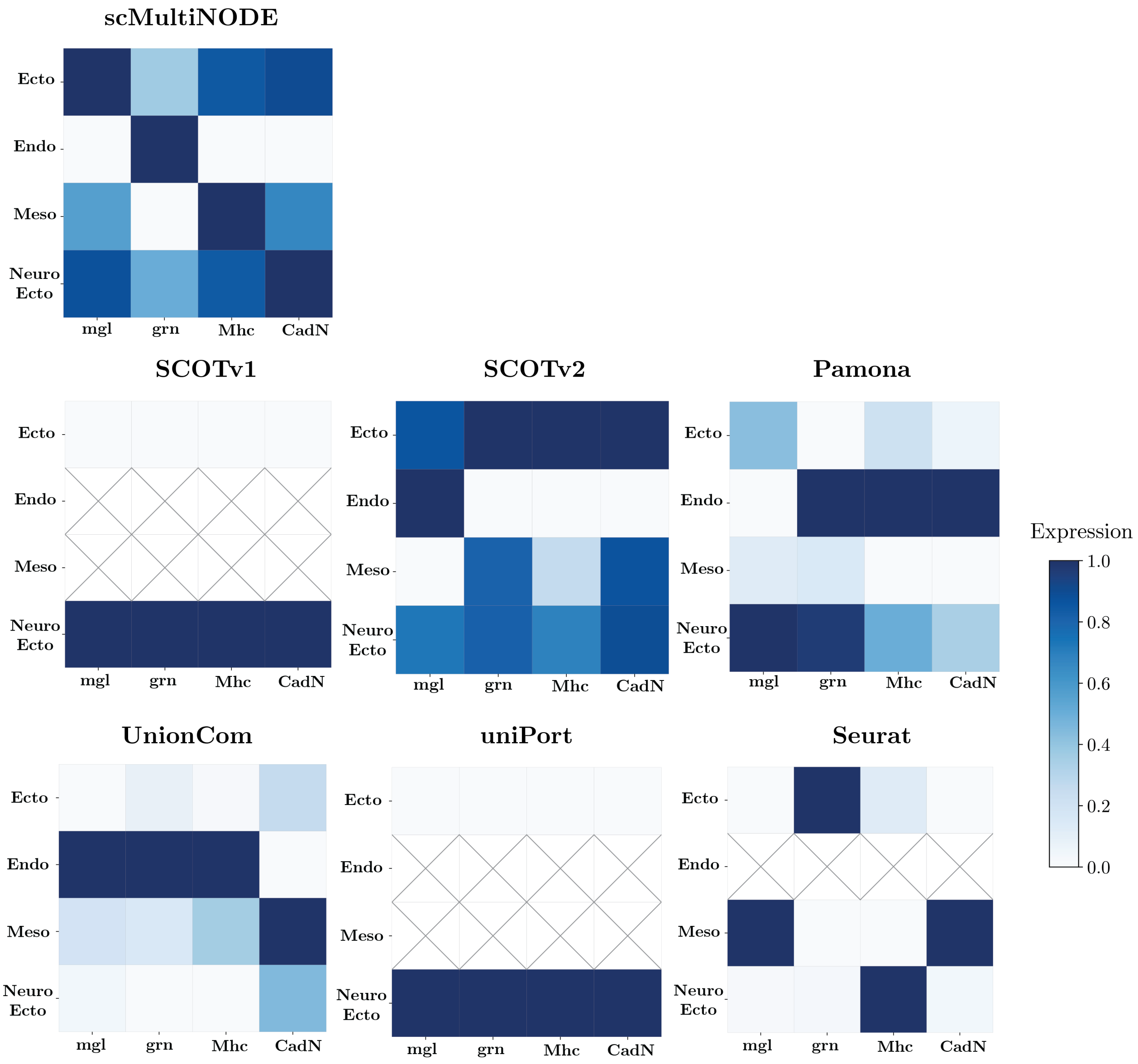


Fig. S13: Expression levels of marker genes (mgl, grn, Mhc, and CadN) across the four predicted germ layer groups (Ecto: ectoderm, Endo: endoderm, Meso: mesoderm, Neuro Ecto: neuroestoderm) from baseline methods' integration. Gene expression values are scaled across all groups. "×" indicates that the classifier fails to predict the corresponding germ layer label based on the integration from that method. We expect to observe a diagonal enrichment pattern, where marker genes are highly expressed in their corresponding cell groups.

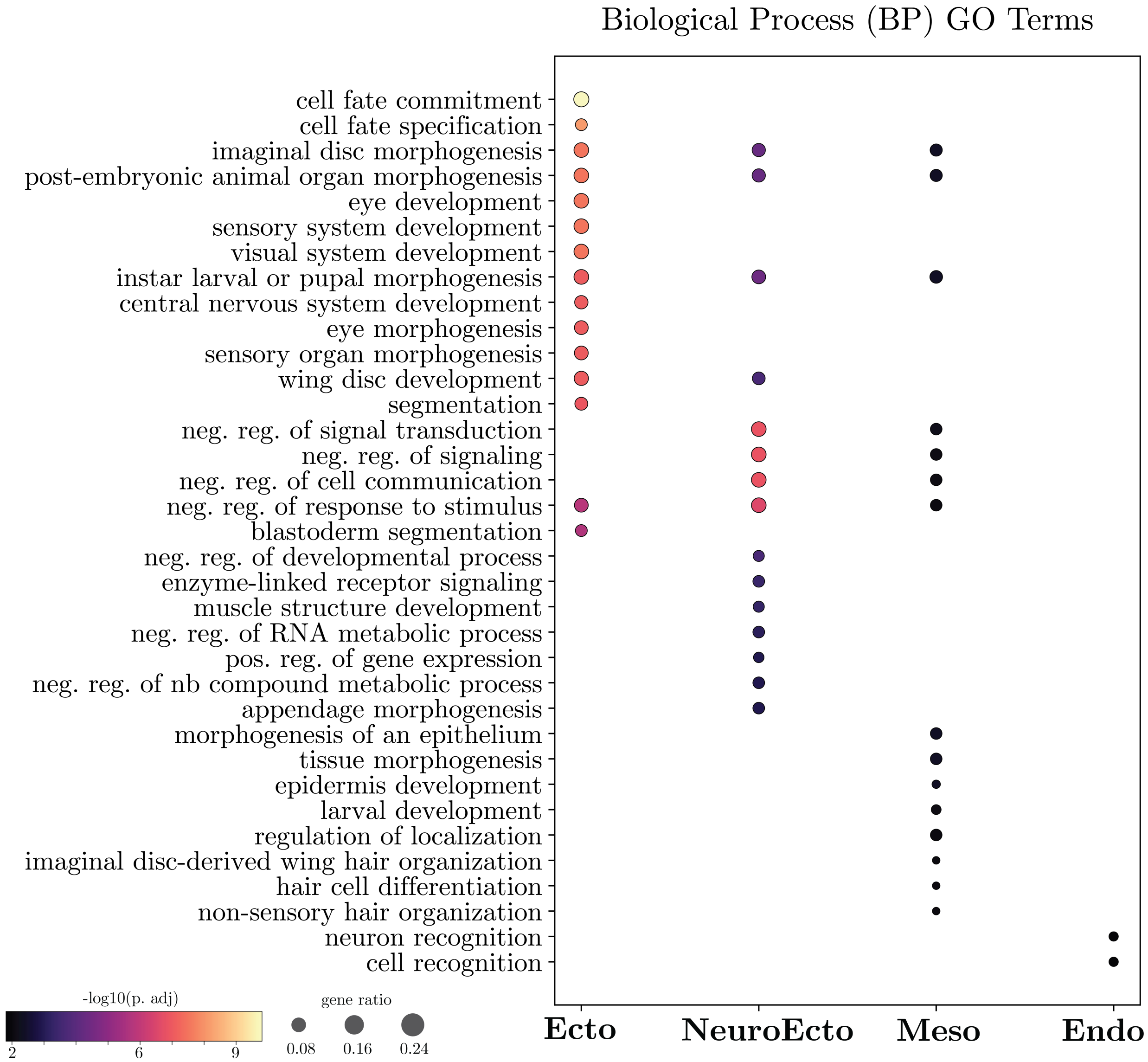


Fig. S14: Gene Ontology (GO) enrichment analysis of the four predicted germ layer groups of `scMultiNODE` , focusing on the Biological Process (BP) category.

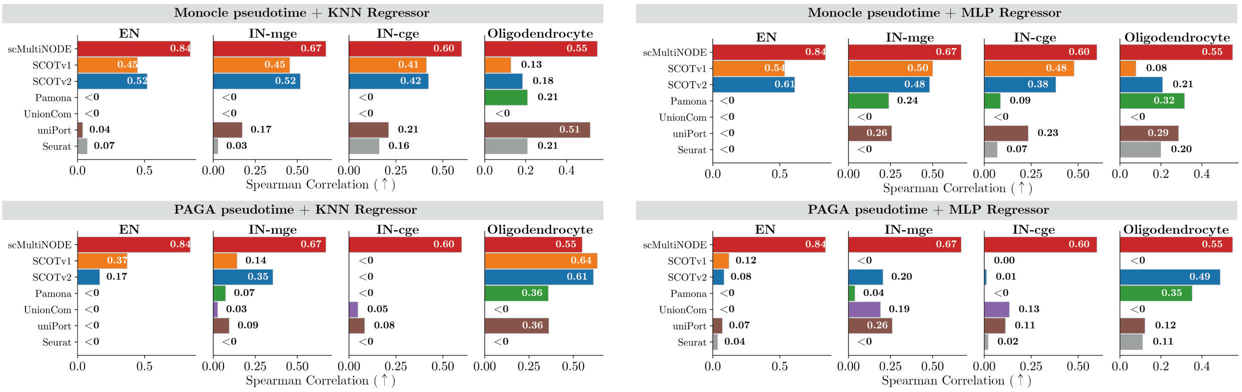


Fig. S15: Performance of `scNODE (transfer)` using different integration methods and regression models. Comparison of pseudotime transfer performance based on Monocle and PAGA pseudotime labels using integrated embeddings from baseline integration methods. `scNODE` is trained on one modality, and the inferred pseudotimes are transferred to the other modality via either a k-nearest neighbors (KNN) regressor or a multilayer perceptron (MLP). Results are averaged over both transfer directions (RNA to ATAC and ATAC to RNA).
